# Improved Identification of Host Cell Proteins in Monoclonal Antibodies by Combining Filter-Aided Sample Preparation and Native Digestion

**DOI:** 10.1101/2025.01.06.631568

**Authors:** Yang Yang, Jing Wu, Fang Wang, Mark Lefers

## Abstract

Host cell proteins (HCPs) in biotherapeutics can present potential safety risks or compromise product stability at trace levels. Therefore, removal, testing, and characterization of HCPs are critical throughout biotherapeutic process development. While enzyme-linked immunosorbent assay (ELISA) is the gold standard for quantifying HCPs, it does not provide information on HCP identity. As a result, HCP analysis by liquid chromatography-mass spectrometry (LC-MS) has gained prominence for its ability to identifying HCPs. However, compared to ELISA’s parts-per-billion sensitivity, LC-MS is limited by its dynamic range, often unable to cover magnitude difference between HCP and biotherapeutic concentrations. Thus, an effective HCP enrichment method is highly desirable. In this study, we propose a new strategy for HCP identification that combines filter-aided sample preparation (FASP) with native digestion, followed by shotgun proteomics analysis. This approach improves mAb removal compared to standard native digestion, enabling greater HCP enrichment and identification. Our method detects all spiked proteins at 1 ppm and most at 0.5 ppm, ranging from 12 to 470 kDa, demonstrating broad molecular weight coverage of HCPs. A proof-of-concept analysis using the NISTmAb standard demonstrates that our filter-aided native digestion identifies 155 more HCPs than the standard native digestion. When applying this strategy to an in-house antibody with low amounts of HCPs, we can quantify HCPs at levels as low as 0.03 ppm, demonstrating the high sensitivity in HCP characterization.

## Introduction

Host cell proteins (HCPs) are process-related impurities found during the production of biotherapeutics in cell lines, such as the Chinese Hamster Ovary (CHO) system. Despite multiple purification steps, trace amounts of HCPs remain in the final drug substance, typically less than 100 parts per million (ppm). These residual HCPs pose several safety and product stability risks: they can exhibit enzymatic activities like cathepsin D, which may bind to monoclonal antibodies (mAbs) and cause fragmentation;^1–3^ they can degrade stabilizer reagents in the formulation buffer, such as esterase family (*e*.*g*., lipase);^4,5^ and they may even trigger immunogenic responses in patients (*e*.*g*., phospholipase B-like 2).^6^ Therefore, it is critical and mandatory to monitor and minimize HCPs throughout the process development to ensure their removal to an acceptable level.^7^

To support high-throughput screening of HCPs, the enzyme-linked immunosorbent assay (ELISA) remains the primary method for quantification, offering high sensitivity down to parts per billion (ppb). However, ELISA only provides a total amount of HCPs, lacking detailed identity information. This limitation particularly weakens its capability for monitoring high-risk HCPs^8^ that require individual identification and quantification. As a result, liquid chromatography-mass spectrometry (LC-MS) has emerged as a powerful alternative, capable of identifying and quantifying HCPs simultaneously. The common sample preparation process in shotgun proteomics includes protein denaturation, disulfide bond reduction, cysteine alkylation, and enzymatic digestion. Although this workflow generates extensive lists of HCP identities, many HCPs remain undetected due to the significant concentration difference between mAbs and HCPs – often as great as six orders of magnitude – exceeding the dynamic range of many modern mass spectrometers. To bridge this large concentration range and enhance sensitivity, several strategies have been developed.^9^ These include HCP enrichment techniques such as concentrating HCPs directly or depleting high concentrations of mAbs through methods like the activity-based protein profiling probe,^4,10^ ProteoMiner enrichment,^10^ affinity chromatography,^11–13^ hydrophilic interaction chromatography (HILIC),^14^ size exclusion chromatography (SEC),^15,16^ molecular weight cutoff (MWCO) separation,^17^ and limited digestion under native conditions.^18^ Additionally, improvements in solution-phase or gas-phase separation methods, such as two-dimensional liquid chromatography (2D LC)^19–21^ or the introduction of ion mobility,^22,23^ have been beneficial. Advanced data acquisition modes like automated iterative^24^ and BoxCar^25^ further facilitate detailed HCP characterization. Together, these innovations have significantly improved HCP detection and characterization, providing crucial insights into the purification process and enhancing overall product safety and stability.

Among these methods, the native digestion (ND) approach^18^ is increasingly being used and is becoming a basis for the development of other related methods for identifying and quantifying HCPs in mAbs. In this study, we propose a simple strategy based on native digestion by integrating filter-aided sample preparation (FASP) into the workflow.^26,27^ By carrying out native digestion on the filter (NDF) instead of in a tube, we have identified significantly more HCPs, resulting from better removal of mAbs without the need of heating and precipitation of the mAb. In contrast to the previous MWCO fractionation method,^17^ which achieves high sensitivity for smaller HCPs but unavoidably loses those larger than the 50 kDa MWCO size, our NDF method offers an unbiased analysis for all spiked proteins ranging from 12 to 470 kDa. This approach enables a more comprehensive HCP profiling and complements the existing platform methods.

### Experimental Section

#### Materials

All chemicals were commercially available and high-purity grade. Chromatography mobile phases (LC-MS grade) were purchased from Thermo Fisher Scientific (Waltham, MA). Monoclonal antibodies (mAb1 & mAb2) were produced by Alexion (New Haven, CT). The NIST monoclonal antibody standard RM 8671 (NISTmAb) was from the National Institute of Standards and Technology (Gaithersburg, MD). Six protein standards were purchased from different vendors including cytochrome c (Cyt c), myoglobin (Mb), phosphorylase b (Phos b), dimeric angiotensin converting enzyme-2 (ACE2),^28^ and fibronectin (FN) were from Sigma-Aldrich (St. Louis, MO); lipoprotein lipase (LPL) was from Bio-Techne (Minneapolis, MN). Two internal standards, bovine β-lactoglobulin and yeast enolase, were obtained from Sigma-Aldrich. Trypsin Platinum (mass spectrometry grade) was purchased from Promega (Madison, WI). The ammonium bicarbonate powder and tris(2-carboxyethyl)phosphine (TCEP) solution with neutral pH were purchased from Thermo Fisher Scientific. Amicon Ultra-0.5, 10 kDa and 3 kDa filters were purchased from MilliporeSigma (Burlington, MA).

#### Comparison between native digestion on the filter (NDF) and native digestion in the tube (ND)

To evaluate the method sensitivity, 10 mg/mL mAb1 with 10 ppm spiked proteins was diluted with non-spiked mAb1 to make samples with 1 and 0.5 ppm spiked proteins. Both NDF and ND underwent buffer exchange of the mAb with spiked protein standards to 50 mM ammonium bicarbonate solution at pH 8.0 three times in a pre-rinsed Amicon Ultra-0.5, 10 kDa filter at 14,000 g for 10 minutes. The ND protocol was adapted from Huang et al.^18^ Briefly, 200 µL of 10 mg/mL mAb with spiked proteins and trypsin at a 1:2,000 enzyme/substrate ratio (w/w) was incubated at 37 °C overnight, followed by disulfide reduction with 2 µL of 500 mM TCEP and mAb precipitation for 10 minutes at 95 °C to enrich HCPs in the supernatant. For the NDF steps, the volume was adjusted to 300 µL of 10 mg/mL mAb with spiked protein standards after buffer exchange. Subsequently, trypsin was added to a 1:2,000 enzyme/substrate ratio (w/w), followed by overnight incubation at 37 °C. The NDF sample was centrifuged at 14,000 g for 15 minutes at room temperature to collect the mAb-depleted solution in the bottom tube. Then, 200 µL of the solution was transferred to a new 1.5 mL Eppendorf tube, ensuring both NDF and ND had the same starting material, 2 mg mAb with HCPs, for direct comparison. To the NDF sample, 2 µL of 500 mM TCEP was added, followed by incubation at 37 °C for 30 minutes. After cooling to room temperature, 5 µL of 10% formic acid was added to both samples, which were then dried in a SpeedVac, reconstituted with 50 µL of 0.1% formic acid. Finally, after measurement with a Nanodrop, 2 µg of the digest of both NDF and ND were injected for LC-MS/MS analysis.

#### LC-MS/MS analysis

All LC-MS/MS analyses were carried out using an Exploris 240 Orbitrap mass spectrometer (Thermo Fisher Scientific, Waltham, MA) connected to a Vanquish Neo UHPLC system (Thermo Fisher Scientific) equipped with an EASY-Spray source. A Trap-and-Elute configuration was employed to inject the digest onto a 5 µm, 300 μm x 5 mm PepMap Neo trap cartridge (Thermo Fisher Scientific), followed by separation on an EASY-Spray PepMap Neo column (100 Å, 2 µm, 75 µm x 500 mm) from Thermo Fisher Scientific. The mobile phases A and B were 0.1% formic acid-based water and acetonitrile, respectively. The flow rate was 300 nL/min, and the gradient increased mobile phase B from 2 to 35% over 160 min, followed by 35 to 80% over 2 min, and maintained at 80% over 13 min. The mass spectrometer used top 20 scans to trigger data-dependent MS2 and other parameters were: MS1 resolution 60,000, MS1 scan range 300-1650 m/z, MS1 AGC target 300%, MS1 maximum injection time 100 ms; MS2 resolution 15,000, MS2 AGC target 100%, MS2 maximum injection time 150 ms. HCD collision energy was 30% and dynamic exclusion was 20 s.

#### Data analysis

The sequences of the in-house mAbs, NISTmAb, spiked proteins, cell line database, and common contaminant proteins were imported into Thermo Proteome Discoverer software 3.1 using the Sequest search engine. The precursor mass tolerance was set at 10 ppm, and the fragment mass tolerance was set at 0.02 Da. Dynamic modifications such as oxidation (+ 15.995 Da), deamidation (+ 0.984 Da), and protein N-terminal acetylation (+ 42.011 Da) were included. The digestion was set to specific “Trypsin (Full)” with a maximum of two missed cleavage sites. The minimal required peptide length was set to six amino acids. The false discovery rate was set at 0.01, and the protein identification was considered to be of high confidence when at least 2 unique peptides were detected. The peak areas from the top 3 peptides (or 2, if only 2 peptides were identified) for each HCP and the areas for two internal standards, bovine β-lactoglobulin and yeast enolase, were used to calculate the concentration of each HCP.

## Results and Discussion

### Effective mAb removal and unbiased HCP detection are achieved through NDF filtration

One pioneering study^17^ first employs a 50 kDa MWCO filter to separate 150 kDa mAbs from HCPs, subsequently digesting the HCP proteins collected in the bottom tube. This method provides high sensitivity, detecting low molecular weight proteins, specifically proteins of 11 and 17 kDa, down to 1 ppm. While this technique effectively meets the detection needs for most HCPs, which are typically smaller than mAbs,^29^ it inadvertently excludes larger HCPs unable to pass through the 50 kDa filter. These include semaphorin-4B (91.4 kDa), sulfhydryl oxidase 1 (82.8 kDa), and polypeptide N-acetylgalactosaminyltransferase 6 (71.9 kDa), thereby posing challenges for comprehensive large HCP analysis. Consequently, a more inclusive HCP profiling approach is desirable.

In our study, we propose a new workflow in which native digestion is performed using a V-shaped MWCO filter to achieve an unbiased analysis, regardless of the size of HCPs. During native digestion on the filter (**Figure 1**), antibodies remain in their intact form due to multiple disulfide linkages, while HCPs are cleaved into peptides. Following digestion, centrifuge filtration separates intact monoclonal antibodies at the top, while HCP peptides pass through the membrane to the bottom for enrichment, irrespective of their size, thereby obtaining a more comprehensive list of HCPs.

**Figure 1.**
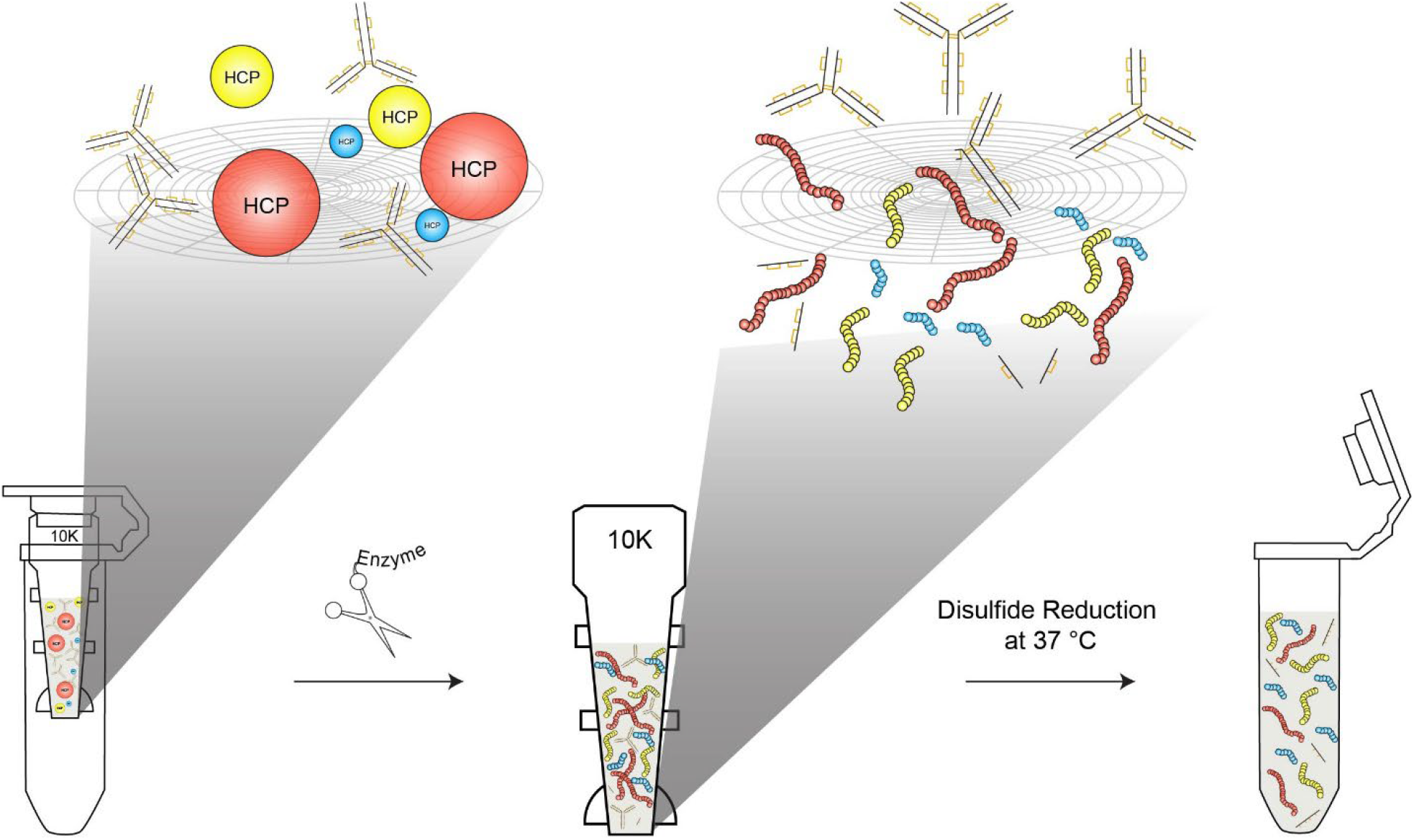
The native digestion on the filter (NDF) experimental workflow for HCP identification includes filter-aided sample preparation and native digestion. During native digestion on the filter, mAbs remain intact due to disulfide bonds, while HCPs are cleaved into peptides. Subsequent centrifuge filtration separates the intact mAbs from the HCP peptides, facilitating HCP enrichment for comprehensive identification, regardless of peptide size.

As process-related samples usually contain salts and other buffer additives (*e*.*g*., 150 mM sodium chloride and polysorbate), which are not compatible with mass spectrometry analysis, a buffer exchange to a volatile salt solution becomes necessary. Therefore, we opt for a 10 kDa filter, which provides the advantage of serving as both a buffer exchanger and digestion reactor simultaneously. Initially, following NDF and centrifuge separation, addition of TCEP and heating at 95 °C for 10 minutes is performed for both NDF and ND, to assess the effectiveness of mAb removal in NDF. In comparison to ND, the NDF digestion recovery tube appears clearer without any cloudiness (**Figure S1A** in the ***Supporting Information***). To confirm the presence of any particulates in NDF, both NDF and ND are centrifuged for 8 minutes. As depicted in **Figure S1B**, NDF remains clear without any visible mAb precipitate, while ND exhibits a significant mAb pellet. This comparison confirms the filter significantly removes the majority of the mAb. In addition, this highlights a unique advantage of NDF in that the high temperature precipitation step can be eliminated, thereby mitigating the risk of loss of heat-labile HCPs or HCPs that get pulled down with the precipitated denatured mAbs. Recent research has shown that after heating and digestion of the pellet in the native digestion process, a significant number of HCPs can further be identified,^30^ which may result from HCP co-precipitation. A plausible hypothesis is that heating inactivates HCPs, such as polysorbate-degrading enzymes, by causing them to unfold and subsequently aggregate,^31^ leading to their precipitation out of the solution. This suggests that developing a method to bypass the heating step is desirable. When comparing NDF with and without the heat precipitation step, the spiked proteins are identified in both conditions, but NDF without heating detects 2 – 3 times more unique peptides for half of the spikes (**Figure S2** in the ***Supporting Information***). Therefore, skipping the heating step not only streamlines the workflow but, more importantly, retains heat-labile species and enables more HCP identifications.

Unlike the previous study,^17^ which uses MWCO enrichment before digestion, the NDF results show that all six protein spikes ranging from 12 kDa to 470 kDa at a 1 ppm level are detected (**Figure 2**), demonstrating no bias towards smaller HCPs and detection of larger HCPs, thereby enabling an unbiased analysis. Among these, the 185 kDa ACE2 exists as a non-covalent dimer.^28^ Its identification further demonstrates that NDF is effective in capturing not only monomeric HCPs but also HCP oligomers in their native form. Importantly, when using the same injection amounts, ND can only detect 2 protein spikes below 20 kDa and loses sensitivity for the other four proteins larger than 50 kDa. This suggests that MWCO filtration may preserve more peptides without heat precipitation, thereby enabling better enrichment of HCPs. Notably, although a 1 ppm sensitivity can be reached for all spikes, the molecular weight of FN is 470 kDa, which is higher than most HCPs or even the high molecular weight impurity (trimeric form) of mAb. This indicates that at least 0.00001 pmol/µL molar concentration of HCP can be confidently detected, demonstrating the satisfactory sensitivity of NDF. When further lowering the spike level down to 0.5 ppm, NDF maintains good detection for all spikes except the largest 470 kDa standard. Although two smaller spikes are still captured by both NDF and ND, the unique peptides reported by NDF are double that of ND, providing more identification confidence and establishing the reliability of the NDF approach.

**Figure 2.**
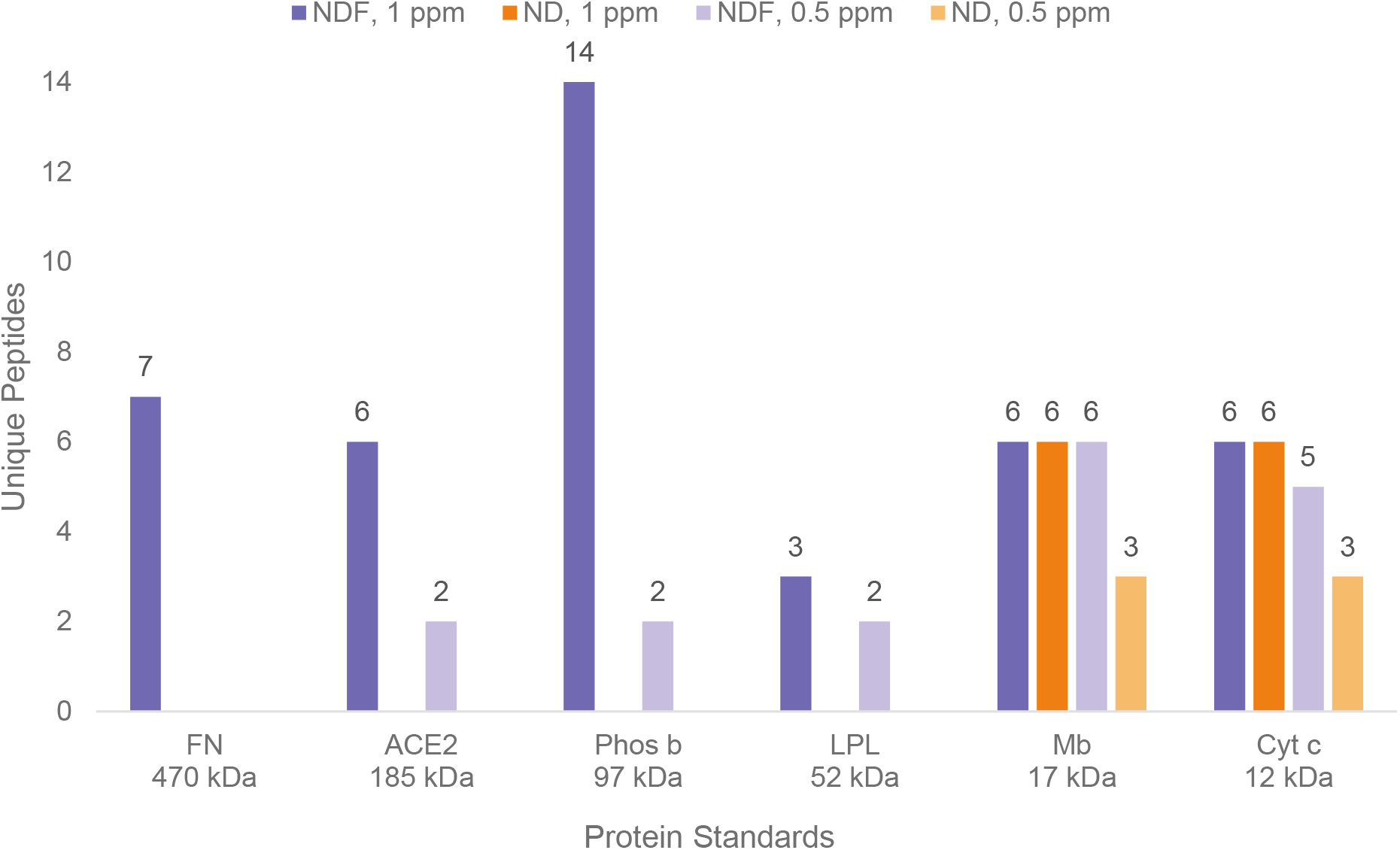
The NDF experiment detects all six protein spikes, ranging from 12 kDa to 470 kDa at 1 ppm, with no bias towards smaller HCPs, allowing for an unbiased analysis. In contrast, ND only detects the Mb and Cyt c spikes, which are below 20 kDa, losing detection for proteins larger than 50 kDa. At a 0.5 ppm spike level, NDF can still detect all spikes except the largest 470 kDa FN and provides nearly twice the unique peptide identification for Mb and Cyt c compared to ND, demonstrating a lower detection limit and offering greater identification confidence in the NDF method.

While the 10 kDa MWCO effectively blocks the mAb on the membrane, the mAb’s signal still dominates and suppresses the recognition of HCP. Considering the link between the mAb removal and the MWCO size, we also examined a 3 kDa MWCO filter. Upon testing the six spiked proteins at 1 ppm, the 3 kDa filter loses detection for half of the protein spikes (**Figure S3** in the ***Supporting Information***), which may indicate that the smaller MWCO size can block larger peptides on the membrane and compromise HCP detection. Therefore, the 10 kDa filter is preferred for the subsequent analysis.

### Identification of HCPs in NISTmAb and a biotherapeutic product by NDF

As the NISTmAb is commonly used as a pharmaceutical standard for method development, we first select the NISTmAb standard (RM 8671) as a case study for direct comparison between NDF and ND sample preparations. As shown in **Figure 3**, both the ND and NDF methods identify 212 HCPs; however, the NDF method detects an additional 165 HCPs, with molecular weights ranging from 11 kDa to 351 kDa (**Table S1** in the ***Supporting Information***). Of these, 160 HCPs can be quantified, with 97% at levels below 5 ppm, 56% below 1 ppm, and one HCP as low as 0.05 ppm (50 ppb). This sensitivity aligns closely with that of ELISA’s ppb range and is also comparable to a targeted analysis with a quantitation limit of 0.06 ppm,^10^ demonstrating the unbiased and sensitive strength of the NDF method. In summary, the NDF method detects an additional 165 HCPs, marking 155 more HCPs than traditional native digestion. In terms of peptides, NDF detects 1,955 unique peptides compared to 1,109 unique peptides detected by ND, indicating that NDF significantly enhances HCP identification by 70% at the protein level and 76% at the peptide level. Therefore, NDF is a reliable method for analyzing low-abundance HCPs with high confidence.

**Figure 3.**
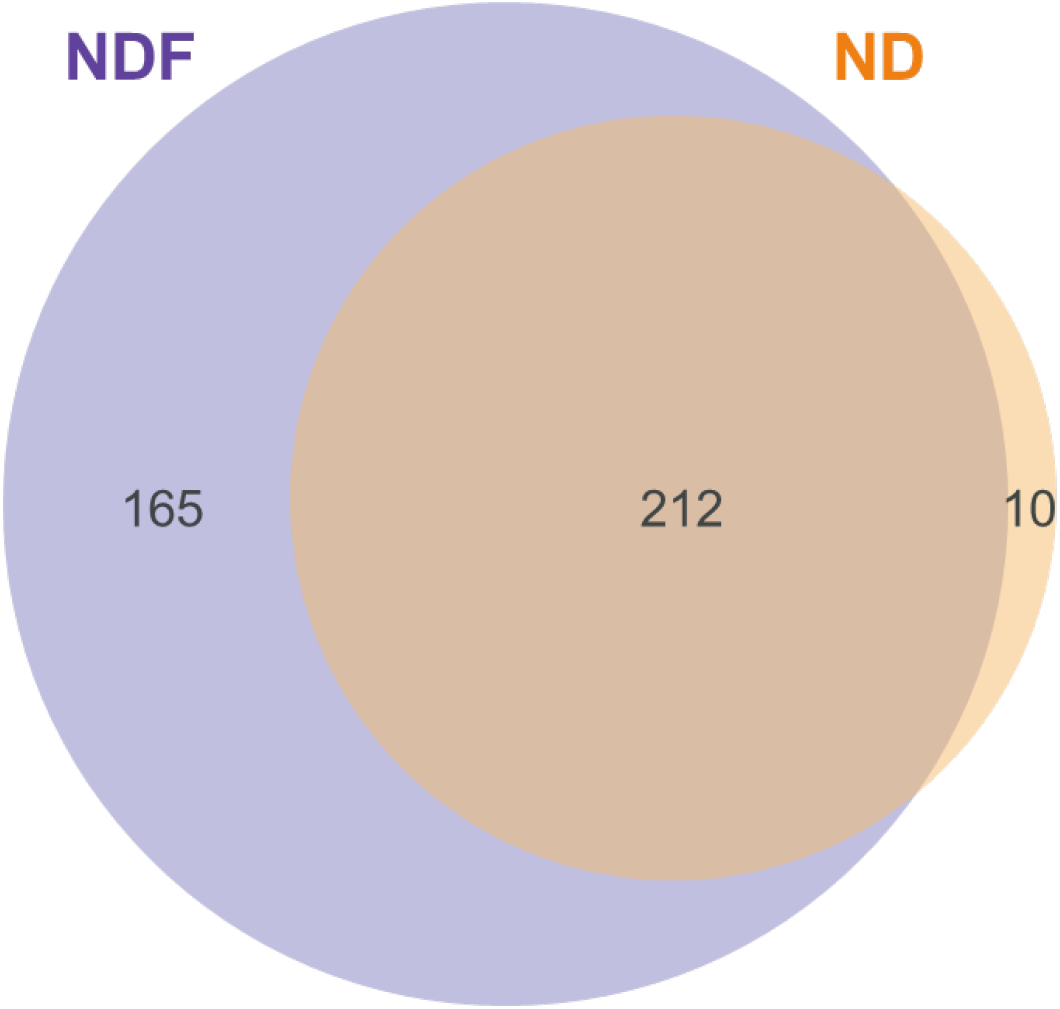
The Venn diagram of total HCPs identified in NISTmAb shows that NDF detects substantially more unique HCPs than ND.

Since the primary objective of conducting HCP analysis in the biopharmaceutical industry is to ensure the suitability of antibody therapeutics, we analyze a high-purity drug substance, mAb2, that undergoes multiple purification steps and has HCP levels significantly below the specified limits. As depicted in **Figure S4** in the ***Supporting Information***, the NDF method for two technical replicates detects an additional 11 HCPs, all quantified at or below 0.2 ppm, with 6 of them at or below 0.1 ppm. The quantification limit reached as low as 0.03 ppm for a 18 kDa myeloid-derived growth factor. Among these 11 HCPs, 6 are classified as high-risk, such as peroxiredoxin-1^32^ at 0.2 ppm and cathepsin D^33^ at 0.1 ppm. These results demonstrate that NDF provides lower limits of quantitation than ND alone, helping the purification department in developing mitigation strategies to continuously enhance the safety and quality of the drug product.

## Conclusions

HCPs are critical quality attributes in the protein drug lifecycle and must be monitored during process development. Consequently, LC-MS/MS is gaining increasing attention as an orthogonal tool to ELISA. However, the limited dynamic range of mass spectrometers poses a challenge for characterizing low-abundance HCPs. Several methods have been introduced at different stages to address this issue, including advancements in data acquisition, second-dimensional separation, and effective HCP enrichment. Among these approaches, native digestion has been more widely adopted owing to its simple setup and robust detection limits.

In our study, we further develop the native digestion strategy by integrating it with filter-aided sample preparation. We demonstrate that NDF is highly effective in removing mAb, while enabling the native digestion of HCPs into peptides and subsequent enrichment in the bottom tube without heating. One recent study^34^ also suggests that intact trypsin remains bound to the filter in the FASP process. Consequently, compared to ND digestion, intact trypsin is negligible in the NDF solution after centrifugation, further reducing the chances of mAb being digested by trypsin, and minimizing signal interference from mAb peptides during HCP detection. When compared to standard native digestion, NDF digestion can achieve a sensitivity of 1 ppm for all and 0.5 ppm for most protein spikes, while standard native digestion has limited capability. In one case study involving NISTmAb, the NDF method detects 155 additional HCPs, with a quantitation limit as low as 0.05 ppm. Notably, this quantitation limit is comparable to a previous study^10^ incorporating parallel reaction monitoring. Such targeted MS/MS analysis is a highly sensitive approach for HCP quantitation; however, it can only be applied when the HCPs of interest are already known, allowing for the selection of specific peptides to monitor routinely. By using the NDF method, similar sensitivity can be achieved during initial HCP profiling, streamlining the workflow and potentially reducing the effort required to develop a targeted method. Moreover, compared to the previous MWCO study,^17^ our method allows for unbiased analysis regardless of HCP size. Given that our method focuses on the initial sample preparation stage, a compelling application would be to integrate NDF with earlier HCP enrichment techniques such as ProteoMiner,^35^ as well as subsequent separation and data acquisition methods, such as 2D LC^20^ and BoxCar scanning,^25^ to further enhance sensitivity.

## Supporting Information

Effective mAb removal by NDF; improved protein and peptide identification by NDF without heating; fewer protein spikes and unique peptides identified by the 3 kDa filter; the Venn diagram of the total HCPs of an in-house drug substance identified by NDF and ND; and total HCPs detected in the NISTmAb by NDF and ND, and unique HCPs detected by NDF.

## Author Information

### Author Contributions

Y.Y. designed the study; Y.Y., J.W. and F.W. designed the experimental work; Y.Y., J.W. and F.W. carried out the experimental work; J.W. and F.W. interpreted the data; Y.Y. and M.L. wrote the manuscript. All authors read the final draft of the manuscript and gave their consent to submitting it for publication.

## Notes

The authors declare no competing financial interest.

## Acknowledgements

This study was sponsored by Alexion, AstraZeneca Rare Disease Unit. The authors are grateful to Dr. Yali Lu (Analytical Sciences, AstraZeneca) for the great discussion and suggestions.

## Table of Contents Graphic

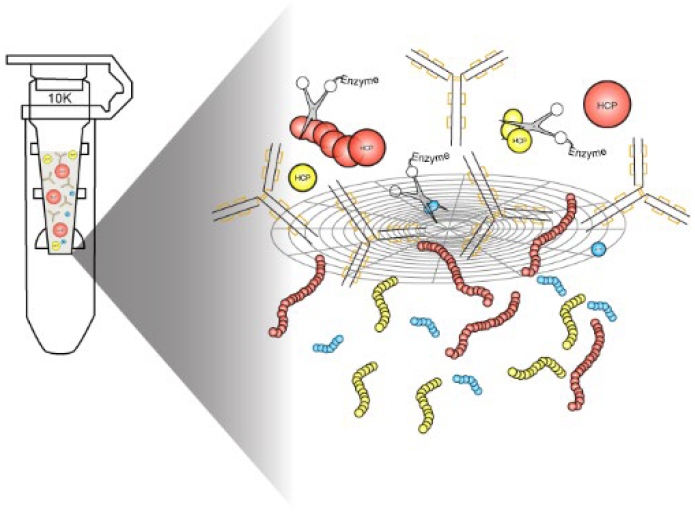

