## Supporting Information for "Improved Identification of Host Cell Proteins in Monoclonal Antibodies by Combining Filter-Aided Sample Preparation and Native Digestion"

*Alexion, AstraZeneca Rare Disease, New Haven, CT*

**Contents**

**Figure S1.** NDF demonstrates effective mAb removal after 95 °C heating, before (**A**) and after (**B**) centrifugation.

**Figure S2.** NDF without 95 °C heating identifies more proteins and peptides, indicating that skipping the heating step retains heat-labile species, reduces HCP co-precipitation, and enables more HCP identifications.

**Figure S3.** The 3 kDa filter identifies fewer protein spikes and unique peptides, which may indicate that the smaller MWCO size can block larger peptides on the membrane and compromise HCP detection.

**Figure S4.** The Venn diagram of the total HCPs of an in-house drug substance identified by NDF and ND.

**Table S1.** Total HCPs detected in the NISTmAb by NDF (**A**) and ND (**B**), and unique HCPs detected by NDF (**C**).

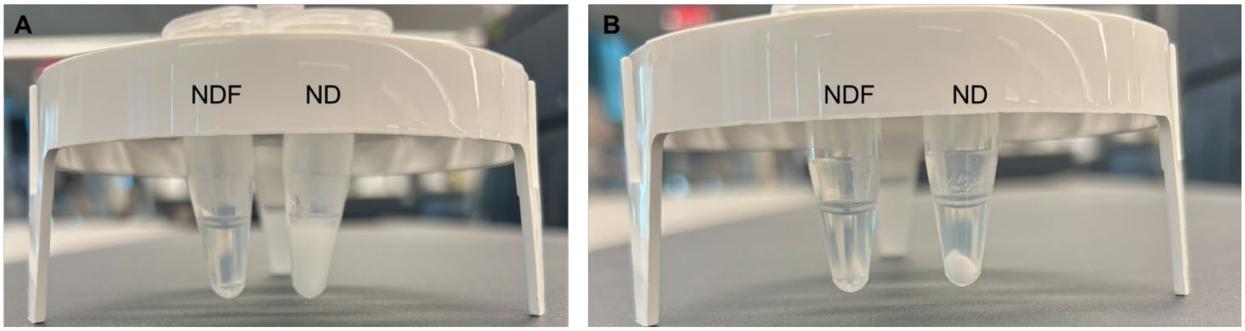


**Figure S1.** Following NDF and centrifuge separation, TCEP is added to both NDF and ND samples at 95 °C for 10 minutes to evaluate mAb removal effectiveness. The NDF tube appears clear compared to the cloudy ND sample (**A**). After centrifugation, NDF remains clear, while ND shows significant mAb presence (**B**), confirming the filter's effective cleanup and high recovery. A key advantage of NDF is the ability to skip high-temperature precipitation, reducing the risk of losing heat-labile HCPs.

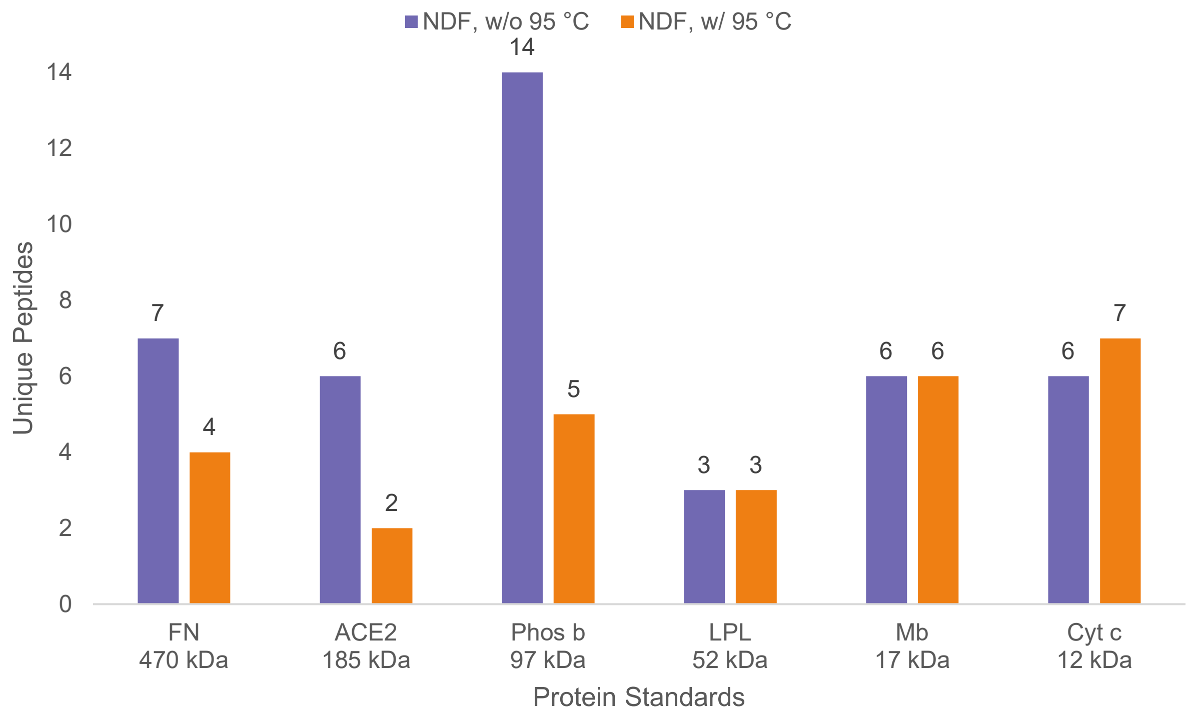


**Figure S2.** Comparison of NDF with and without the heat precipitation step shows that while both conditions identify spiked proteins, NDF without heating detects 2 – 3 times more unique peptides for half of the spikes, such as FN, ACE2, and Phos b. This implies that skipping the heating step may enhance the retention of heat-labile species or reduce the risk of HCPs being co-precipitated with denatured mAbs, enabling more comprehensive HCP identifications.

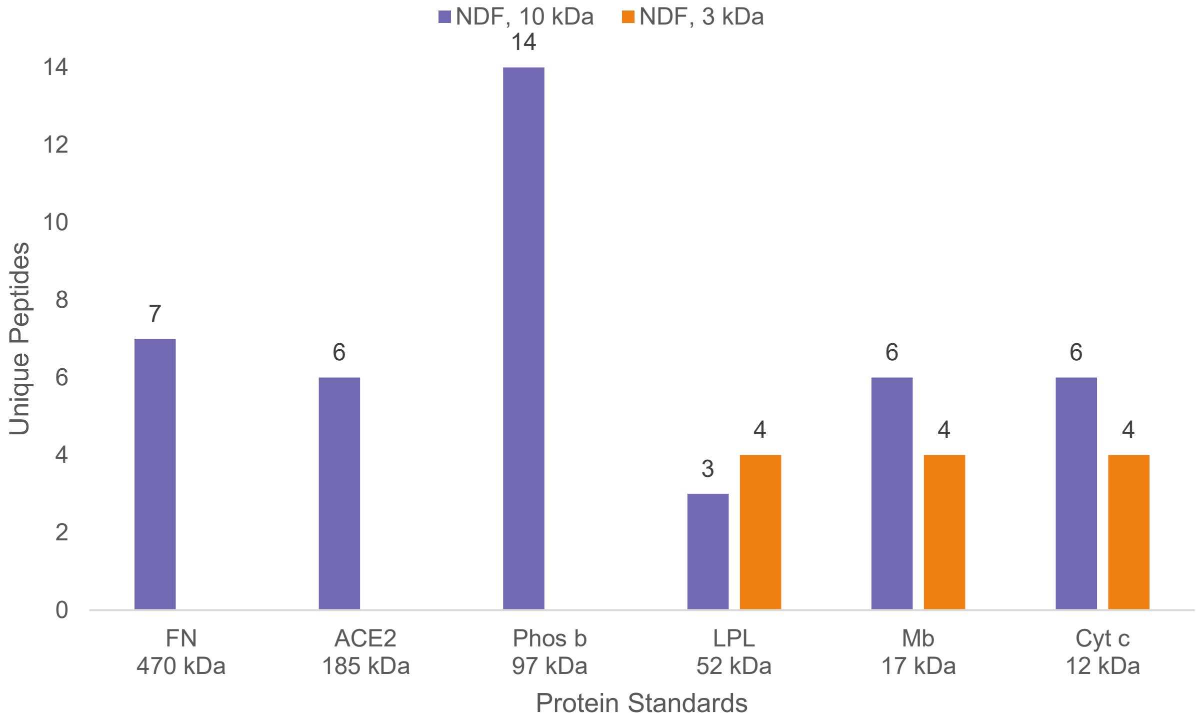


**Figure S3.** Examination of a 3 kDa filter (catalogue no. UFC500324) on mAb blockage shows that half of the protein spikes are unidentified at 1 ppm, including 470 kDa FN, 185 kDa ACE2, and 97 kDa Phos b, and that 17 kDa Mb and 12 kDa Cyt c have fewer peptides detected than the 10 kDa filter (catalogue no. UFC501024). This suggests that a smaller MWCO size may block larger peptides, thereby compromising HCP detection.

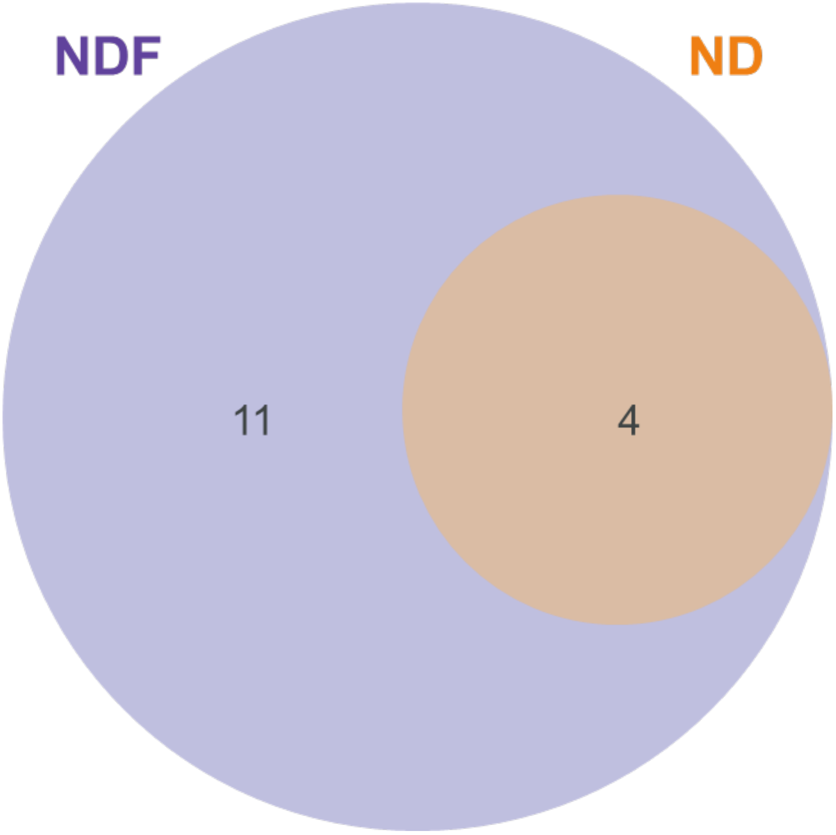


**Figure S4.** The Venn diagram of total HCPs identified in an in-house drug substance shows that NDF detects significantly more unique HCPs than ND.

**Table S1.** Across two technical replicates for HCP identification in NISTmAb, the NDF method identifies 377 CHO proteins (**A**) in contrast to 222 HCPs detected by ND (**B**). NDF uniquely detects an additional 165 HCPs (**C**) with molecular weights from 11 kDa to 351 kDa, demonstrating high sensitivity across a broad mass range.

(**A**)

| Accession | Description | MW, kDa | # Unique Peptides |
|---|---|---|---|
| P62858 | Small ribosomal subunit protein eS28 OS=Mus musculus OX=10090 GN=Rps28 PE=1 SV=1 | 7.8 | 2 |
| P62075 | Mitochondrial import inner membrane translocase subunit Tim13 OS=Mus musculus OX=10090 GN=Timm13 PE=1 SV=1 | 10.5 | 4 |
| Q64433 | 10 kDa heat shock protein, mitochondrial OS=Mus musculus OX=10090 GN=Hspe1 PE=1 SV=2 | 11.0 | 3 |
| Q9JIX0 | Transcription and mRNA export factor ENY2 OS=Mus musculus OX=10090 GN=Eny2 PE=1 SV=1 | 11.5 | 5 |
| P62897 | Cytochrome c, somatic OS=Mus musculus OX=10090 GN=Cycs PE=1 SV=2 | 11.6 | 5 |
| O35143 | ATPase inhibitor, mitochondrial OS=Mus musculus OX=10090 GN=Atp5if1 PE=1 SV=2 | 12.2 | 5 |
| P83870 | PHD finger-like domain-containing protein 5A OS=Mus musculus OX=10090 GN=Phf5a PE=1 SV=1 | 12.4 | 3 |
| Q9CWW6 | Peptidyl-prolyl cis-trans isomerase NIMA-interacting 4 OS=Mus musculus OX=10090 GN=Pin4 PE=1 SV=1 | 13.8 | 3 |
| P70349 | Adenosine 5'-monophosphoramidase HINT1 OS=Mus musculus OX=10090 GN=Hint1 PE=1 SV=3 | 13.8 | 2 |
| P01887 | Beta-2-microglobulin OS=Mus musculus OX=10090 GN=B2m PE=1 SV=2 | 13.8 | 4 |
| P56812 | Programmed cell death protein 5 OS=Mus musculus OX=10090 GN=Pdcd5 PE=1 SV=3 | 14.3 | 6 |
| Q9D8S9 | BolA-like protein 1 OS=Mus musculus OX=10090 GN=Bola1 PE=1 SV=1 | 14.4 | 2 |
| Q03958 | Prefoldin subunit 6 OS=Mus musculus OX=10090 GN=Pfdn6 PE=1 SV=1 | 14.4 | 6 |
| P63323 | Small ribosomal subunit protein eS12 OS=Mus musculus OX=10090 GN=Rps12 PE=1 SV=3 | 14.5 | 6 |
| P59708 | Splicing factor 3B subunit 6 OS=Mus musculus OX=10090 GN=Sf3b6 PE=1 SV=1 | 14.6 | 2 |
| P45878 | Peptidyl-prolyl cis-trans isomerase FKBP2 OS=Mus musculus OX=10090 GN=Fkbp2 PE=1 SV=1 | 15.3 | 9 |
| P21460 | Cystatin-C OS=Mus musculus OX=10090 GN=Cst3 PE=1 SV=2 | 15.5 | 5 |
| Q9CR98 | Protein FAM136A OS=Mus musculus OX=10090 GN=Fam136a PE=1 SV=1 | 15.7 | 2 |
| Q9CZG9 | PDZ domain-containing protein 11 OS=Mus musculus OX=10090 GN=Pdzd11 PE=1 SV=1 | 16.2 | 2 |
| Q8VHC3 | Selenoprotein M OS=Mus musculus OX=10090 GN=Selenom PE=1 SV=3 | 16.4 | 5 |
| Q9D2M8 | Ubiquitin-conjugating enzyme E2 variant 2 OS=Mus musculus OX=10090 GN=Ube2v2 PE=1 SV=4 | 16.4 | 3 |
| O70591 | Prefoldin subunit 2 OS=Mus musculus OX=10090 GN=Pfdn2 PE=1 SV=2 | 16.5 | 2 |
| Q8R3Q6 | Protein MIX23 OS=Mus musculus OX=10090 GN=Mix23 PE=1 SV=1 | 16.7 | 4 |

| | | | |
|---|---|---|---|
| O70200 | Allograft inflammatory factor 1 OS=Mus musculus OX=10090 GN=Aif1 PE=1 SV=1 | 16.9 | 4 |
| Q9CQ92 | Mitochondrial fission 1 protein OS=Mus musculus OX=10090 GN=Fis1 PE=1 SV=1 | 17.0 | 3 |
| P62843 | Small ribosomal subunit protein uS19 OS=Mus musculus OX=10090 GN=Rps15 PE=1 SV=2 | 17.0 | 3 |
| P61089 | Ubiquitin-conjugating enzyme E2 N OS=Mus musculus OX=10090 GN=Ube2n PE=1 SV=1 | 17.1 | 2 |
| P54227 | Stathmin OS=Mus musculus OX=10090 GN=Stmn1 PE=1 SV=2 | 17.3 | 3 |
| Q01768 | Nucleoside diphosphate kinase B OS=Mus musculus OX=10090 GN=Nme2 PE=1 SV=1 | 17.4 | 2 |
| P62270 | Small ribosomal subunit protein uS13 OS=Mus musculus OX=10090 GN=Rps18 PE=1 SV=3 | 17.7 | 4 |
| P21126 | Ubiquitin-like protein 4A OS=Mus musculus OX=10090 GN=Ubl4a PE=1 SV=1 | 17.8 | 2 |
| P62983 | Ubiquitin-ribosomal protein eS31 fusion protein OS=Mus musculus OX=10090 GN=Rps27a PE=1 SV=2 | 17.9 | 6 |
| Q9DCT6 | Chromatin complexes subunit BAP18 OS=Mus musculus OX=10090 GN=Bap18 PE=1 SV=1 | 18.0 | 2 |
| P17742 | Peptidyl-prolyl cis-trans isomerase A OS=Mus musculus OX=10090 GN=Ppia PE=1 SV=2 | 18.0 | 4 |
| Q9D7P6 | Iron-sulfur cluster assembly enzyme ISCU OS=Mus musculus OX=10090 GN=Iscu PE=1 SV=1 | 18.1 | 8 |
| P18760 | Cofilin-1 OS=Mus musculus OX=10090 GN=Cfl1 PE=1 SV=3 | 18.5 | 3 |
| Q8R2Q8 | Bone marrow stromal antigen 2 OS=Mus musculus OX=10090 GN=Bst2 PE=1 SV=1 | 19.1 | 4 |
| P53996 | CCHC-type zinc finger nucleic acid binding protein OS=Mus musculus OX=10090 GN=Cnbp PE=1 SV=2 | 19.6 | 2 |
| Q9CXZ1 | NADH dehydrogenase [ubiquinone] iron-sulfur protein 4, mitochondrial OS=Mus musculus OX=10090 GN=Ndufs4 PE=1 SV=3 | 19.8 | 2 |
| Q9D1M4 | Eukaryotic translation elongation factor 1 epsilon-1 OS=Mus musculus OX=10090 GN=Eef1e1 PE=1 SV=1 | 19.8 | 6 |
| Q6PGH2 | Jupiter microtubule associated homolog 2 OS=Mus musculus OX=10090 GN=Jpt2 PE=1 SV=1 | 20.0 | 3 |
| Q99LX0 | Parkinson disease protein 7 homolog OS=Mus musculus OX=10090 GN=Park7 PE=1 SV=1 | 20.0 | 5 |
| P61205 | ADP-ribosylation factor 3 OS=Mus musculus OX=10090 GN=Arf3 PE=2 SV=2 | 20.6 | 3 |
| Q9DB15 | Large ribosomal subunit protein bL12m OS=Mus musculus OX=10090 GN=Mrpl12 PE=1 SV=2 | 21.7 | 3 |
| Q99KR7 | Peptidyl-prolyl cis-trans isomerase F, mitochondrial OS=Mus musculus OX=10090 GN=Ppif PE=1 SV=1 | 21.7 | 3 |
| O88952 | Protein lin-7 homolog C OS=Mus musculus OX=10090 GN=Lin7c PE=1 SV=2 | 21.8 | 4 |
| Q61171 | Peroxiredoxin-2 OS=Mus musculus OX=10090 GN=Prdx2 PE=1 SV=3 | 21.8 | 3 |
| P99029 | Peroxiredoxin-5, mitochondrial OS=Mus musculus OX=10090 GN=Prdx5 PE=1 SV=2 | 21.9 | 8 |
| P35700 | Peroxiredoxin-1 OS=Mus musculus OX=10090 GN=Prdx1 PE=1 SV=1 | 22.2 | 12 |
| Q9DBP5 | UMP-CMP kinase OS=Mus musculus OX=10090 GN=Cmpk1 PE=1 SV=1 | 22.2 | 5 |

| | | | |
|---|---|---|---|
| Q923D2 | Flavin reductase (NADPH) OS=Mus musculus OX=10090 GN=Blvrb PE=1 SV=3 | 22.2 | 9 |
| Q9CQW1 | Synaptobrevin homolog YKT6 OS=Mus musculus OX=10090 GN=Ykt6 PE=1 SV=1 | 22.3 | 3 |
| Q9WVA4 | Transgelin-2 OS=Mus musculus OX=10090 GN=Tagln2 PE=1 SV=4 | 22.4 | 12 |
| Q8VCG4 | Complement component C8 gamma chain OS=Mus musculus OX=10090 GN=C8g PE=1 SV=1 | 22.5 | 4 |
| P56213 | FAD-linked sulfhydryl oxidase ALR OS=Mus musculus OX=10090 GN=Gfer PE=1 SV=2 | 22.9 | 2 |
| Q8BPA8 | Protein DPCD OS=Mus musculus OX=10090 GN=Dpcd PE=1 SV=1 | 23.0 | 6 |
| Q9DCT5 | Stromal cell-derived factor 2 OS=Mus musculus OX=10090 GN=Sdf2 PE=1 SV=1 | 23.1 | 2 |
| Q6IRU5-2 | Isoform 2 of Clathrin light chain B OS=Mus musculus OX=10090 GN=Cltb | 23.2 | 7 |
| P19157 | Glutathione S-transferase P 1 OS=Mus musculus OX=10090 GN=Gstp1 PE=1 SV=2 | 23.6 | 6 |
| P24369 | Peptidyl-prolyl cis-trans isomerase B OS=Mus musculus OX=10090 GN=Ppib PE=1 SV=2 | 23.7 | 9 |
| P30681 | High mobility group protein B2 OS=Mus musculus OX=10090 GN=Hmgb2 PE=1 SV=3 | 24.1 | 5 |
| Q78PG9 | Coiled-coil domain-containing protein 25 OS=Mus musculus OX=10090 GN=Ccdc25 PE=1 SV=1 | 24.5 | 3 |
| P09671 | Superoxide dismutase [Mn], mitochondrial OS=Mus musculus OX=10090 GN=Sod2 PE=1 SV=3 | 24.6 | 5 |
| O35166 | Golgi SNAP receptor complex member 2 OS=Mus musculus OX=10090 GN=Gosr2 PE=1 SV=2 | 24.7 | 6 |
| P63158 | High mobility group protein B1 OS=Mus musculus OX=10090 GN=Hmgb1 PE=1 SV=2 | 24.9 | 3 |
| O08709 | Peroxiredoxin-6 OS=Mus musculus OX=10090 GN=Prdx6 PE=1 SV=3 | 24.9 | 10 |
| Q9D8B3 | Charged multivesicular body protein 4b OS=Mus musculus OX=10090 GN=Chmp4b PE=1 SV=2 | 24.9 | 5 |
| Q62446 | Peptidyl-prolyl cis-trans isomerase FKBP3 OS=Mus musculus OX=10090 GN=Fkbp3 PE=1 SV=2 | 25.1 | 4 |
| Q60631 | Growth factor receptor-bound protein 2 OS=Mus musculus OX=10090 GN=Grb2 PE=1 SV=1 | 25.2 | 6 |
| Q9ERE7 | LRP chaperone MESD OS=Mus musculus OX=10090 GN=Mesd PE=1 SV=1 | 25.2 | 2 |
| Q62093 | Serine/arginine-rich splicing factor 2 OS=Mus musculus OX=10090 GN=Srsf2 PE=1 SV=4 | 25.5 | 3 |
| Q61206 | Platelet-activating factor acetylhydrolase IB subunit alpha2 OS=Mus musculus OX=10090 GN=Pafah1b2 PE=1 SV=2 | 25.6 | 3 |
| Q61205 | Platelet-activating factor acetylhydrolase IB subunit alpha1 OS=Mus musculus OX=10090 GN=Pafah1b3 PE=1 SV=1 | 25.8 | 6 |
| P10649 | Glutathione S-transferase Mu 1 OS=Mus musculus OX=10090 GN=Gstm1 PE=1 SV=2 | 26.0 | 2 |
| Q9CQF3 | Cleavage and polyadenylation specificity factor subunit 5 OS=Mus musculus OX=10090 GN=Nudt21 PE=1 SV=1 | 26.2 | 7 |
| Q9CRB9 | MICOS complex subunit Mic19 OS=Mus musculus OX=10090 GN=Chchd3 PE=1 SV=1 | 26.3 | 2 |
| Q8C7V8 | Coiled-coil domain-containing protein 134 OS=Mus musculus OX=10090 GN=Ccdc134 PE=1 SV=1 | 26.5 | 3 |

| | | | |
|---|---|---|---|
| Q9WTP6 | Adenylate kinase 2, mitochondrial OS=Mus musculus OX=10090 GN=Ak2 PE=1 SV=5 | 26.5 | 14 |
| Q9CXW3 | Calcyclin-binding protein OS=Mus musculus OX=10090 GN=Cacybp PE=1 SV=1 | 26.5 | 2 |
| P17751 | Triosephosphate isomerase OS=Mus musculus OX=10090 GN=Tpi1 PE=1 SV=5 | 26.7 | 2 |
| O88983 | Syntaxin-8 OS=Mus musculus OX=10090 GN=Stx8 PE=1 SV=1 | 26.9 | 2 |
| O08583 | THO complex subunit 4 OS=Mus musculus OX=10090 GN=Alyref PE=1 SV=3 | 26.9 | 3 |
| Q03402 | Cysteine-rich secretory protein 3 OS=Mus musculus OX=10090 GN=Crisp3 PE=1 SV=1 | 27.3 | 4 |
| O09131 | Glutathione S-transferase omega-1 OS=Mus musculus OX=10090 GN=Gsto1 PE=1 SV=2 | 27.5 | 5 |
| Q6PDM2 | Serine/arginine-rich splicing factor 1 OS=Mus musculus OX=10090 GN=Srsf1 PE=1 SV=3 | 27.7 | 6 |
| Q9WV55 | Vesicle-associated membrane protein-associated protein A OS=Mus musculus OX=10090 GN=Vapa PE=1 SV=2 | 27.8 | 4 |
| Q61335 | B-cell receptor-associated protein 31 OS=Mus musculus OX=10090 GN=Bcap31 PE=1 SV=4 | 27.9 | 3 |
| Q9CQE8 | RNA transcription, translation and transport factor protein OS=Mus musculus OX=10090 GN=RTRAF PE=1 SV=1 | 28.1 | 4 |
| Q9D172 | Glutamine amidotransferase-like class 1 domain-containing protein 3, mitochondrial OS=Mus musculus OX=10090 GN=Gatd3 PE=1 SV=1 | 28.1 | 4 |
| Q9CQE1 | Protein NipSnap homolog 3B OS=Mus musculus OX=10090 GN=Nipsnap3b PE=1 SV=1 | 28.3 | 7 |
| Q9D1J1 | Adaptin ear-binding coat-associated protein 2 OS=Mus musculus OX=10090 GN=Necap2 PE=1 SV=1 | 28.6 | 2 |
| O70250 | Phosphoglycerate mutase 2 OS=Mus musculus OX=10090 GN=Pgam2 PE=1 SV=3 | 28.8 | 4 |
| P62259 | 14-3-3 protein epsilon OS=Mus musculus OX=10090 GN=Ywhae PE=1 SV=1 | 29.2 | 2 |
| Q99020 | Heterogeneous nuclear ribonucleoprotein A/B OS=Mus musculus OX=10090 GN=Hnrnpab PE=1 SV=1 | 30.8 | 7 |
| Q8BL97 | Serine/arginine-rich splicing factor 7 OS=Mus musculus OX=10090 GN=Srsf7 PE=1 SV=1 | 30.8 | 7 |
| O35326 | Serine/arginine-rich splicing factor 5 OS=Mus musculus OX=10090 GN=Srsf5 PE=1 SV=2 | 30.9 | 3 |
| Q9ER00 | Syntaxin-12 OS=Mus musculus OX=10090 GN=Stx12 PE=1 SV=1 | 31.2 | 8 |
| Q9R0Q4 | Mortality factor 4-like protein 2 OS=Mus musculus OX=10090 GN=Morf4l2 PE=1 SV=1 | 32.2 | 5 |
| Q61937 | Nucleophosmin OS=Mus musculus OX=10090 GN=Npm1 PE=1 SV=1 | 32.5 | 4 |
| Q9QZH3 | Peptidyl-prolyl cis-trans isomerase E OS=Mus musculus OX=10090 GN=Ppie PE=1 SV=2 | 33.4 | 2 |
| Q8K4F5 | sn-1-specific diacylglycerol lipase ABHD11 OS=Mus musculus OX=10090 GN=Abhd11 PE=1 SV=1 | 33.5 | 10 |
| Q922Y1 | UBX domain-containing protein 1 OS=Mus musculus OX=10090 GN=Ubxn1 PE=1 SV=1 | 33.6 | 4 |
| P10711 | Transcription elongation factor A protein 1 OS=Mus musculus OX=10090 GN=Tcea1 PE=1 SV=2 | 33.9 | 2 |
| Q8C0M9 | Isoaspartyl peptidase/L-asparaginase OS=Mus musculus OX=10090 GN=Asrgl1 PE=1 SV=1 | 33.9 | 7 |

| | | | |
|---|---|---|---|
| Q9WUU7 | Cathepsin Z OS=Mus musculus OX=10090 GN=Ctsz PE=1 SV=1 | 34.0 | 2 |
| P31230 | Aminoacyl tRNA synthase complex-interacting multifunctional protein 1 OS=Mus musculus OX=10090 GN=Aimp1 PE=1 SV=2 | 34.0 | 4 |
| Q99KB8 | Hydroxyacylglutathione hydrolase, mitochondrial OS=Mus musculus OX=10090 GN=Hagh PE=1 SV=2 | 34.1 | 2 |
| P49312 | Heterogeneous nuclear ribonucleoprotein A1 OS=Mus musculus OX=10090 GN=Hnrnpa1 PE=1 SV=2 | 34.2 | 10 |
| Q9Z204 | Heterogeneous nuclear ribonucleoproteins C1/C2 OS=Mus musculus OX=10090 GN=Hnrnpc PE=1 SV=1 | 34.4 | 5 |
| Q61425 | Hydroxyacyl-coenzyme A dehydrogenase, mitochondrial OS=Mus musculus OX=10090 GN=Hadh PE=1 SV=2 | 34.4 | 5 |
| P56528 | ADP-ribosyl cyclase/cyclic ADP-ribose hydrolase 1 OS=Mus musculus OX=10090 GN=Cd38 PE=1 SV=2 | 34.4 | 2 |
| O88531 | Palmitoyl-protein thioesterase 1 OS=Mus musculus OX=10090 GN=Ppt1 PE=1 SV=2 | 34.5 | 5 |
| Q8BM88 | Cathepsin O OS=Mus musculus OX=10090 GN=Ctso PE=2 SV=1 | 34.7 | 2 |
| Q61176 | Arginase-1 OS=Mus musculus OX=10090 GN=Arg1 PE=1 SV=1 | 34.8 | 4 |
| Q91Z53 | Glyoxylate reductase/hydroxypyruvate reductase OS=Mus musculus OX=10090 GN=Grhpr PE=1 SV=1 | 35.3 | 2 |
| Q91YR9 | Prostaglandin reductase 1 OS=Mus musculus OX=10090 GN=Ptgr1 PE=1 SV=2 | 35.5 | 14 |
| Q9Z1D1 | Eukaryotic translation initiation factor 3 subunit G OS=Mus musculus OX=10090 GN=Eif3g PE=1 SV=2 | 35.6 | 2 |
| P08249 | Malate dehydrogenase, mitochondrial OS=Mus musculus OX=10090 GN=Mdh2 PE=1 SV=3 | 35.6 | 11 |
| P45376 | Aldo-keto reductase family 1 member B1 OS=Mus musculus OX=10090 GN=Akr1b1 PE=1 SV=3 | 35.7 | 5 |
| P62960 | Y-box-binding protein 1 OS=Mus musculus OX=10090 GN=Ybx1 PE=1 SV=3 | 35.7 | 3 |
| P16858 | Glyceraldehyde-3-phosphate dehydrogenase OS=Mus musculus OX=10090 GN=Gapdh PE=1 SV=2 | 35.8 | 2 |
| P70372 | ELAV-like protein 1 OS=Mus musculus OX=10090 GN=Elavl1 PE=1 SV=2 | 36.1 | 9 |
| P14152 | Malate dehydrogenase, cytoplasmic OS=Mus musculus OX=10090 GN=Mdh1 PE=1 SV=3 | 36.5 | 2 |
| P06151 | L-lactate dehydrogenase A chain OS=Mus musculus OX=10090 GN=Ldha PE=1 SV=3 | 36.5 | 9 |
| P08101 | Low affinity immunoglobulin gamma Fc region receptor II OS=Mus musculus OX=10090 GN=Fcgr2 PE=1 SV=2 | 36.7 | 3 |
| Q9D1P4 | Cysteine and histidine-rich domain-containing protein 1 OS=Mus musculus OX=10090 GN=Chordc1 PE=1 SV=1 | 37.3 | 2 |
| Q93092 | Transaldolase OS=Mus musculus OX=10090 GN=Taldo1 PE=1 SV=2 | 37.4 | 12 |
| Q9ES89 | Exostosin-like 2 OS=Mus musculus OX=10090 GN=Extl2 PE=1 SV=1 | 37.4 | 6 |
| O88569 | Heterogeneous nuclear ribonucleoproteins A2/B1 OS=Mus musculus OX=10090 GN=Hnrnpa2b1 PE=1 SV=2 | 37.4 | 9 |
| P06797 | Procathepsin L OS=Mus musculus OX=10090 GN=Ctsl PE=1 SV=2 | 37.5 | 4 |
| P60335 | Poly(rC)-binding protein 1 OS=Mus musculus OX=10090 GN=Pcbp1 PE=1 SV=1 | 37.5 | 5 |

| | | | |
|---|---|---|---|
| Q8VCN9 | Tubulin-specific chaperone C OS=Mus musculus OX=10090 GN=Tbcc PE=1 SV=1 | 38.1 | 2 |
| Q64442 | Sorbitol dehydrogenase OS=Mus musculus OX=10090 GN=Sord PE=1 SV=3 | 38.2 | 6 |
| O35685 | Nuclear migration protein nudC OS=Mus musculus OX=10090 GN=Nudc PE=1 SV=1 | 38.3 | 2 |
| P70441 | Na(+)/H(+) exchange regulatory cofactor NHE-RF1 OS=Mus musculus OX=10090 GN=Nherf1 PE=1 SV=3 | 38.6 | 5 |
| P56542 | Deoxyribonuclease-2-alpha OS=Mus musculus OX=10090 GN=Dnase2 PE=1 SV=1 | 38.8 | 9 |
| P24452 | Macrophage-capping protein OS=Mus musculus OX=10090 GN=Capg PE=1 SV=2 | 39.2 | 4 |
| Q3UMW8 | Bis(monoacylglycero)phosphate synthase CLN5 OS=Mus musculus OX=10090 GN=Cln5 PE=1 SV=1 | 39.3 | 3 |
| P05064 | Fructose-bisphosphate aldolase A OS=Mus musculus OX=10090 GN=Aldoa PE=1 SV=2 | 39.3 | 30 |
| P05063 | Fructose-bisphosphate aldolase C OS=Mus musculus OX=10090 GN=Aldoc PE=1 SV=4 | 39.4 | 24 |
| Q9JHJ0 | Tropomodulin-3 OS=Mus musculus OX=10090 GN=Tmod3 PE=1 SV=1 | 39.5 | 2 |
| P28474 | Alcohol dehydrogenase class-3 OS=Mus musculus OX=10090 GN=Adh5 PE=1 SV=3 | 39.5 | 5 |
| Q8BG05 | Heterogeneous nuclear ribonucleoprotein A3 OS=Mus musculus OX=10090 GN=Hnrnpa3 PE=1 SV=1 | 39.6 | 2 |
| P54726 | UV excision repair protein RAD23 homolog A OS=Mus musculus OX=10090 GN=Rad23a PE=1 SV=2 | 39.7 | 2 |
| O54946 | DnaJ homolog subfamily B member 6 OS=Mus musculus OX=10090 GN=Dnajb6 PE=1 SV=4 | 39.8 | 4 |
| Q9CX56 | 26S proteasome non-ATPase regulatory subunit 8 OS=Mus musculus OX=10090 GN=Psmd8 PE=1 SV=2 | 39.9 | 3 |
| P59481 | VIP36-like protein OS=Mus musculus OX=10090 GN=Lman2l PE=1 SV=1 | 39.9 | 5 |
| Q78JW9 | Ubiquitin domain-containing protein UBFD1 OS=Mus musculus OX=10090 GN=Ubfd1 PE=1 SV=2 | 40.1 | 5 |
| Q91YR1 | Twinfilin-1 OS=Mus musculus OX=10090 GN=Twf1 PE=1 SV=2 | 40.1 | 4 |
| Q00731-6 | Isoform L-VEGF-1 of Vascular endothelial growth factor A, long form OS=Mus musculus OX=10090 GN=Vegfa | 40.3 | 2 |
| Q9CR16 | Peptidyl-prolyl cis-trans isomerase D OS=Mus musculus OX=10090 GN=Ppid PE=1 SV=3 | 40.7 | 10 |
| Q9CZ44 | NSFL1 cofactor p47 OS=Mus musculus OX=10090 GN=Nsfl1c PE=1 SV=1 | 40.7 | 21 |
| P63085 | Mitogen-activated protein kinase 1 OS=Mus musculus OX=10090 GN=Mapk1 PE=1 SV=3 | 41.2 | 4 |
| Q8CAY6 | Acetyl-CoA acetyltransferase, cytosolic OS=Mus musculus OX=10090 GN=Acat2 PE=1 SV=2 | 41.3 | 4 |
| Q9Z0P4 | Paralemmin-1 OS=Mus musculus OX=10090 GN=Palm PE=1 SV=1 | 41.6 | 2 |
| Q91VM5 | RNA binding motif protein, X-linked-like-1 OS=Mus musculus OX=10090 GN=Rbmxl1 PE=2 SV=1 | 42.1 | 4 |
| P34902 | Cytokine receptor common subunit gamma OS=Mus musculus OX=10090 GN=Il2rg PE=1 SV=1 | 42.2 | 5 |
| P55302 | Alpha-2-macroglobulin receptor-associated protein OS=Mus musculus OX=10090 GN=Lrpap1 PE=1 SV=1 | 42.2 | 8 |

| | | | |
|---|---|---|---|
| P70318 | Nucleolysin TIAR OS=Mus musculus OX=10090 GN=Tial1 PE=1 SV=1 | 43.4 | 3 |
| P50580 | Proliferation-associated protein 2G4 OS=Mus musculus OX=10090 GN=Pa2g4 PE=1 SV=3 | 43.7 | 2 |
| Q61187 | Tumor susceptibility gene 101 protein OS=Mus musculus OX=10090 GN=Tsg101 PE=1 SV=2 | 44.1 | 2 |
| P04202 | Transforming growth factor beta-1 proprotein OS=Mus musculus OX=10090 GN=Tgfb1 PE=1 SV=1 | 44.3 | 5 |
| P15535 | Beta-1,4-galactosyltransferase 1 OS=Mus musculus OX=10090 GN=B4galt1 PE=1 SV=1 | 44.4 | 2 |
| P09411 | Phosphoglycerate kinase 1 OS=Mus musculus OX=10090 GN=Pgk1 PE=1 SV=4 | 44.5 | 10 |
| Q9CY58 | SERPINE1 mRNA-binding protein 1 OS=Mus musculus OX=10090 GN=Serbp1 PE=1 SV=2 | 44.7 | 2 |
| P18242 | Cathepsin D OS=Mus musculus OX=10090 GN=Ctsd PE=1 SV=1 | 44.9 | 9 |
| O89112 | Glutathione S-transferase LANCL1 OS=Mus musculus OX=10090 GN=Lancl1 PE=1 SV=1 | 45.3 | 3 |
| Q8BK62 | Olfactomedin-like protein 3 OS=Mus musculus OX=10090 GN=Olfml3 PE=2 SV=2 | 45.7 | 2 |
| Q8BV49 | Pyrin and HIN domain-containing protein 1 OS=Mus musculus OX=10090 GN=Pyhin1 PE=1 SV=1 | 46.9 | 2 |
| Q8BHS3 | Pre-mRNA-splicing factor RBM22 OS=Mus musculus OX=10090 GN=Rbm22 PE=1 SV=1 | 46.9 | 2 |
| P17182 | Alpha-enolase OS=Mus musculus OX=10090 GN=Eno1 PE=1 SV=3 | 47.1 | 8 |
| Q9QWR8 | Alpha-N-acetylgalactosaminidase OS=Mus musculus OX=10090 GN=Naga PE=1 SV=2 | 47.2 | 2 |
| Q8BP40 | Lysophosphatidic acid phosphatase type 6 OS=Mus musculus OX=10090 GN=Acp6 PE=1 SV=1 | 47.6 | 3 |
| P50247 | Adenosylhomocysteinase OS=Mus musculus OX=10090 GN=Ahcy PE=1 SV=3 | 47.7 | 9 |
| Q922R8 | Protein disulfide-isomerase A6 OS=Mus musculus OX=10090 GN=Pdia6 PE=1 SV=3 | 48.1 | 9 |
| Q62418 | Drebrin-like protein OS=Mus musculus OX=10090 GN=Dbnl PE=1 SV=2 | 48.7 | 8 |
| Q8VEJ9 | Vacuolar protein sorting-associated protein 4A OS=Mus musculus OX=10090 GN=Vps4a PE=1 SV=1 | 48.9 | 2 |
| O35737 | Heterogeneous nuclear ribonucleoprotein H OS=Mus musculus OX=10090 GN=Hnrnph1 PE=1 SV=3 | 49.2 | 3 |
| Q9JIY5 | Serine protease HTRA2, mitochondrial OS=Mus musculus OX=10090 GN=Htra2 PE=1 SV=2 | 49.3 | 5 |
| P11680 | Properdin OS=Mus musculus OX=10090 GN=Cfp PE=1 SV=2 | 50.3 | 3 |
| Q80V42 | Carboxypeptidase M OS=Mus musculus OX=10090 GN=Cpm PE=1 SV=2 | 50.5 | 4 |
| O55131 | Septin-7 OS=Mus musculus OX=10090 GN=Septin7 PE=1 SV=1 | 50.5 | 4 |
| Q9QUN3 | B-cell linker protein OS=Mus musculus OX=10090 GN=Blnk PE=1 SV=1 | 50.6 | 4 |
| P61979 | Heterogeneous nuclear ribonucleoprotein K OS=Mus musculus OX=10090 GN=Hnrnpk PE=1 SV=1 | 50.9 | 4 |
| Q64287 | Interferon regulatory factor 4 OS=Mus musculus OX=10090 GN=Irf4 PE=1 SV=1 | 51.5 | 10 |

| | | | |
|---|---|---|---|
| P40124 | Adenylyl cyclase-associated protein 1 OS=Mus musculus OX=10090 GN=Cap1 PE=1 SV=4 | 51.5 | 8 |
| P27808 | Alpha-1,3-mannosyl-glycoprotein 2-beta-N-acetylglucosaminyltransferase OS=Mus musculus OX=10090 GN=Mgat1 PE=1 SV=1 | 51.7 | 2 |
| P97855 | Ras GTPase-activating protein-binding protein 1 OS=Mus musculus OX=10090 GN=G3bp1 PE=1 SV=1 | 51.8 | 5 |
| Q9Z2W0 | Aspartyl aminopeptidase OS=Mus musculus OX=10090 GN=Dnpep PE=1 SV=2 | 52.2 | 3 |
| Q6NXH2 | Glycoprotein endo-alpha-1,2-mannosidase OS=Mus musculus OX=10090 GN=Manea PE=2 SV=1 | 53.1 | 7 |
| Q9DCD0 | 6-phosphogluconate dehydrogenase, decarboxylating OS=Mus musculus OX=10090 GN=Pgd PE=1 SV=3 | 53.2 | 8 |
| Q02819 | Nucleobindin-1 OS=Mus musculus OX=10090 GN=Nucb1 PE=1 SV=2 | 53.4 | 2 |
| Q56A08 | G-patch domain and KOW motifs-containing protein OS=Mus musculus OX=10090 GN=Gpkow PE=1 SV=2 | 53.8 | 3 |
| P17892 | Pancreatic lipase-related protein 2 OS=Mus musculus OX=10090 GN=Pnliprp2 PE=1 SV=2 | 54.0 | 9 |
| Q2TPA8 | Hydroxysteroid dehydrogenase-like protein 2 OS=Mus musculus OX=10090 GN=Hsdl2 PE=1 SV=1 | 54.2 | 2 |
| P49710 | Hematopoietic lineage cell-specific protein OS=Mus musculus OX=10090 GN=Hcls1 PE=1 SV=2 | 54.2 | 3 |
| P97807 | Fumarate hydratase, mitochondrial OS=Mus musculus OX=10090 GN=Fh PE=1 SV=3 | 54.3 | 12 |
| Q99K48 | Non-POU domain-containing octamer-binding protein OS=Mus musculus OX=10090 GN=Nono PE=1 SV=3 | 54.5 | 2 |
| Q02853 | Stromelysin-3 OS=Mus musculus OX=10090 GN=Mmp11 PE=1 SV=2 | 55.4 | 3 |
| Q8BG07 | 5'-3' exonuclease PLD4 OS=Mus musculus OX=10090 GN=Pld4 PE=1 SV=1 | 56.1 | 5 |
| Q9QYI3 | DnaJ homolog subfamily C member 7 OS=Mus musculus OX=10090 GN=Dnajc7 PE=1 SV=2 | 56.4 | 5 |
| P97360 | Transcription factor ETV6 OS=Mus musculus OX=10090 GN=Etv6 PE=1 SV=1 | 56.4 | 4 |
| Q8VCH8 | UBX domain-containing protein 4 OS=Mus musculus OX=10090 GN=Ubxn4 PE=1 SV=1 | 56.4 | 3 |
| Q61753 | D-3-phosphoglycerate dehydrogenase OS=Mus musculus OX=10090 GN=Phgdh PE=1 SV=3 | 56.5 | 4 |
| O35664 | Interferon alpha/beta receptor 2 OS=Mus musculus OX=10090 GN=Ifnar2 PE=1 SV=2 | 56.5 | 3 |
| Q99K28 | ADP-ribosylation factor GTPase-activating protein 2 OS=Mus musculus OX=10090 GN=Arfgap2 PE=1 SV=1 | 56.6 | 5 |
| Q9D8S3 | ADP-ribosylation factor GTPase-activating protein 3 OS=Mus musculus OX=10090 GN=Arfgap3 PE=1 SV=2 | 57.4 | 5 |
| P09242 | Alkaline phosphatase, tissue-nonspecific isozyme OS=Mus musculus OX=10090 GN=Alpl PE=1 SV=2 | 57.5 | 5 |
| Q8BG30 | Negative elongation factor A OS=Mus musculus OX=10090 GN=Nelfa PE=1 SV=1 | 57.5 | 3 |
| P52480 | Pyruvate kinase PKM OS=Mus musculus OX=10090 GN=Pkm PE=1 SV=4 | 57.8 | 5 |
| Q3UEB3-2 | Isoform 2 of Poly(U)-binding-splicing factor PUF60 OS=Mus musculus OX=10090 GN=Puf60 | 58.5 | 13 |
| O08795 | Glucosidase 2 subunit beta OS=Mus musculus OX=10090 GN=Prkcsh PE=1 SV=1 | 58.8 | 2 |

| | | | |
|---|---|---|---|
| Q6NVF9 | Cleavage and polyadenylation specificity factor subunit 6 OS=Mus musculus OX=10090 GN=Cpsf6 PE=1 SV=1 | 59.1 | 5 |
| Q9Z2A5 | Arginyl-tRNA--protein transferase 1 OS=Mus musculus OX=10090 GN=Ate1 PE=1 SV=2 | 59.1 | 3 |
| P32020 | Sterol carrier protein 2 OS=Mus musculus OX=10090 GN=Scp2 PE=1 SV=3 | 59.1 | 9 |
| P17225 | Polypyrimidine tract-binding protein 1 OS=Mus musculus OX=10090 GN=Ptbp1 PE=1 SV=3 | 59.3 | 5 |
| Q8CI11 | Guanine nucleotide-binding protein-like 3 OS=Mus musculus OX=10090 GN=Gnl3 PE=1 SV=2 | 60.7 | 3 |
| P63038 | 60 kDa heat shock protein, mitochondrial OS=Mus musculus OX=10090 GN=Hspd1 PE=1 SV=1 | 60.9 | 4 |
| P20060 | Beta-hexosaminidase subunit beta OS=Mus musculus OX=10090 GN=Hexb PE=1 SV=2 | 61.1 | 11 |
| Q8BFR4 | N-acetylglucosamine-6-sulfatase OS=Mus musculus OX=10090 GN=Gns PE=1 SV=1 | 61.1 | 2 |
| Q8BIQ5 | Cleavage stimulation factor subunit 2 OS=Mus musculus OX=10090 GN=Cstf2 PE=1 SV=2 | 61.3 | 2 |
| Q60864 | Stress-induced-phosphoprotein 1 OS=Mus musculus OX=10090 GN=Stip1 PE=1 SV=1 | 62.5 | 13 |
| P03975 | IgE-binding protein OS=Mus musculus OX=10090 GN=Iap PE=2 SV=1 | 62.7 | 9 |
| P06745 | Glucose-6-phosphate isomerase OS=Mus musculus OX=10090 GN=Gpi PE=1 SV=4 | 62.7 | 23 |
| Q9D2L1 | Arylsulfatase K OS=Mus musculus OX=10090 GN=Arsk PE=1 SV=2 | 62.8 | 8 |
| Q8C854 | Myelin expression factor 2 OS=Mus musculus OX=10090 GN=Myef2 PE=1 SV=1 | 63.3 | 4 |
| Q8CHU3 | Epsin-2 OS=Mus musculus OX=10090 GN=Epn2 PE=1 SV=1 | 63.4 | 7 |
| P28798 | Progranulin OS=Mus musculus OX=10090 GN=Grn PE=1 SV=2 | 63.4 | 4 |
| Q61712 | DnaJ homolog subfamily C member 1 OS=Mus musculus OX=10090 GN=Dnajc1 PE=1 SV=1 | 63.8 | 3 |
| Q80WJ7 | Protein LYRIC OS=Mus musculus OX=10090 GN=Mtdh PE=1 SV=1 | 63.8 | 7 |
| Q6PB93 | Polypeptide N-acetylgalactosaminyltransferase 2 OS=Mus musculus OX=10090 GN=Galnt2 PE=1 SV=1 | 64.5 | 25 |
| Q60862 | Origin recognition complex subunit 2 OS=Mus musculus OX=10090 GN=Orc2 PE=1 SV=1 | 65.9 | 2 |
| Q925I1 | ATPase family AAA domain-containing protein 3 OS=Mus musculus OX=10090 GN=Atad3 PE=1 SV=1 | 66.7 | 2 |
| Q9Z0X1 | Apoptosis-inducing factor 1, mitochondrial OS=Mus musculus OX=10090 GN=Aifm1 PE=1 SV=1 | 66.7 | 18 |
| Q8CHY6 | Transcriptional repressor p66 alpha OS=Mus musculus OX=10090 GN=Gatad2a PE=1 SV=2 | 67.3 | 2 |
| P40142 | Transketolase OS=Mus musculus OX=10090 GN=Tkt PE=1 SV=1 | 67.6 | 26 |
| P35235-1 | Isoform 2 of Tyrosine-protein phosphatase non-receptor type 11 OS=Mus musculus OX=10090 GN=Ptpn11 | 68.4 | 7 |
| Q61545 | RNA-binding protein EWS OS=Mus musculus OX=10090 GN=Ewsr1 PE=1 SV=2 | 68.4 | 4 |
| Q91WJ8 | Far upstream element-binding protein 1 OS=Mus musculus OX=10090 GN=Fubp1 PE=1 SV=1 | 68.5 | 5 |

| | | | |
|---|---|---|---|
| Q99KN9 | Clathrin interactor 1 OS=Mus musculus OX=10090 GN=Clint1 PE=1 SV=2 | 68.5 | 7 |
| Q8BGD9 | Eukaryotic translation initiation factor 4B OS=Mus musculus OX=10090 GN=Eif4b PE=1 SV=1 | 68.8 | 9 |
| P10404 | MLV-related proviral Env polyprotein OS=Mus musculus OX=10090 PE=1 SV=3 | 69.6 | 2 |
| Q9DBG7 | Signal recognition particle receptor subunit alpha OS=Mus musculus OX=10090 GN=Srpra PE=1 SV=1 | 69.6 | 3 |
| Q9JL61 | DNA-binding protein Rfx5 OS=Mus musculus OX=10090 GN=Rfx5 PE=1 SV=2 | 69.7 | 2 |
| Q5RKZ7 | Molybdenum cofactor biosynthesis protein 1 OS=Mus musculus OX=10090 GN=Mocs1 PE=1 SV=2 | 69.8 | 5 |
| Q8R0H9 | ADP-ribosylation factor-binding protein GGA1 OS=Mus musculus OX=10090 GN=Gga1 PE=1 SV=1 | 69.9 | 2 |
| P54729 | NEDD8 ultimate buster 1 OS=Mus musculus OX=10090 GN=Nub1 PE=1 SV=2 | 70.3 | 2 |
| Q64213 | Splicing factor 1 OS=Mus musculus OX=10090 GN=Sf1 PE=1 SV=6 | 70.4 | 4 |
| Q9JLQ0 | CD2-associated protein OS=Mus musculus OX=10090 GN=Cd2ap PE=1 SV=3 | 70.4 | 13 |
| P29341 | Polyadenylate-binding protein 1 OS=Mus musculus OX=10090 GN=Pabpc1 PE=1 SV=2 | 70.6 | 11 |
| P63017 | Heat shock cognate 71 kDa protein OS=Mus musculus OX=10090 GN=Hspa8 PE=1 SV=1 | 70.8 | 13 |
| Q80TY0 | Formin-binding protein 1 OS=Mus musculus OX=10090 GN=Fnbp1 PE=1 SV=2 | 71.3 | 3 |
| Q8K2Q9 | Shootin-1 OS=Mus musculus OX=10090 GN=Shtn1 PE=1 SV=1 | 71.3 | 2 |
| Q3U9G9 | Delta(14)-sterol reductase LBR OS=Mus musculus OX=10090 GN=Lbr PE=1 SV=2 | 71.4 | 5 |
| Q8C7U7 | Polypeptide N-acetylgalactosaminyltransferase 6 OS=Mus musculus OX=10090 GN=Galnt6 PE=2 SV=1 | 71.5 | 25 |
| P54103 | DnaJ homolog subfamily C member 2 OS=Mus musculus OX=10090 GN=Dnajc2 PE=1 SV=2 | 71.7 | 6 |
| P08003 | Protein disulfide-isomerase A4 OS=Mus musculus OX=10090 GN=Pdia4 PE=1 SV=3 | 71.9 | 3 |
| Q8BVL9 | Janus kinase and microtubule-interacting protein 1 OS=Mus musculus OX=10090 GN=Jakmip1 PE=1 SV=2 | 73.1 | 2 |
| Q8VC60 | Beta-galactosidase-1-like protein OS=Mus musculus OX=10090 GN=Glb1l PE=1 SV=1 | 73.2 | 5 |
| Q3V1H1 | Cytoskeleton-associated protein 2 OS=Mus musculus OX=10090 GN=Ckap2 PE=1 SV=1 | 74.0 | 2 |
| Q9D706 | RNA polymerase II-associated protein 3 OS=Mus musculus OX=10090 GN=Rpap3 PE=1 SV=1 | 74.1 | 9 |
| P48678 | Prelamin-A/C OS=Mus musculus OX=10090 GN=Lmna PE=1 SV=2 | 74.2 | 14 |
| Q9QUR8 | Semaphorin-7A OS=Mus musculus OX=10090 GN=Sema7a PE=1 SV=1 | 74.9 | 16 |
| Q8VIJ6 | Splicing factor, proline- and glutamine-rich OS=Mus musculus OX=10090 GN=Sfpq PE=1 SV=1 | 75.4 | 7 |
| Q5F2E7 | FMR1-interacting protein NUFIP2 OS=Mus musculus OX=10090 GN=Nufip2 PE=1 SV=1 | 75.6 | 2 |
| P09405 | Nucleolin OS=Mus musculus OX=10090 GN=Ncl PE=1 SV=2 | 76.7 | 2 |

| | | | |
|---|---|---|---|
| Q3U0V1 | Far upstream element-binding protein 2 OS=Mus musculus OX=10090 GN=Khsrp PE=1 SV=2 | 76.7 | 6 |
| Q9D0E1 | Heterogeneous nuclear ribonucleoprotein M OS=Mus musculus OX=10090 GN=Hnrnpm PE=1 SV=3 | 77.6 | 4 |
| P51660 | Peroxisomal multifunctional enzyme type 2 OS=Mus musculus OX=10090 GN=Hsd17b4 PE=1 SV=3 | 79.4 | 2 |
| Q8BWW4 | La-related protein 4 OS=Mus musculus OX=10090 GN=Larp4 PE=1 SV=2 | 79.7 | 2 |
| O88967 | ATP-dependent zinc metalloprotease YME1L1 OS=Mus musculus OX=10090 GN=Yme1l1 PE=1 SV=1 | 80.0 | 7 |
| P26928 | Hepatocyte growth factor-like protein OS=Mus musculus OX=10090 GN=Mst1 PE=2 SV=2 | 80.6 | 15 |
| Q08943 | FACT complex subunit SSRP1 OS=Mus musculus OX=10090 GN=Ssrp1 PE=1 SV=2 | 80.8 | 2 |
| Q6A0A2 | La-related protein 4B OS=Mus musculus OX=10090 GN=Larp4b PE=1 SV=2 | 81.6 | 8 |
| Q9CZD3 | Glycine--tRNA ligase OS=Mus musculus OX=10090 GN=Gars1 PE=1 SV=1 | 81.8 | 2 |
| Q61183 | Poly(A) polymerase alpha OS=Mus musculus OX=10090 GN=Papola PE=1 SV=4 | 82.3 | 2 |
| Q8BJ05 | Zinc finger CCCH domain-containing protein 14 OS=Mus musculus OX=10090 GN=Zc3h14 PE=1 SV=1 | 82.4 | 4 |
| Q8BND5 | Sulfhydryl oxidase 1 OS=Mus musculus OX=10090 GN=Qsox1 PE=1 SV=1 | 82.7 | 16 |
| Q3UIR3 | E3 ubiquitin-protein ligase DTX3L OS=Mus musculus OX=10090 GN=Dtx3l PE=1 SV=1 | 83.0 | 6 |
| Q9D0R2 | Threonine--tRNA ligase 1, cytoplasmic OS=Mus musculus OX=10090 GN=Tars1 PE=1 SV=2 | 83.3 | 2 |
| P51125 | Calpastatin OS=Mus musculus OX=10090 GN=Cast PE=1 SV=2 | 84.9 | 2 |
| Q68FF6 | ARF GTPase-activating protein GIT1 OS=Mus musculus OX=10090 GN=Git1 PE=1 SV=1 | 85.2 | 5 |
| Q99KI0 | Aconitate hydratase, mitochondrial OS=Mus musculus OX=10090 GN=Aco2 PE=1 SV=1 | 85.4 | 17 |
| Q9QYH6 | Melanoma-associated antigen D1 OS=Mus musculus OX=10090 GN=Maged1 PE=1 SV=1 | 85.6 | 2 |
| Q62351 | Transferrin receptor protein 1 OS=Mus musculus OX=10090 GN=Tfrc PE=1 SV=1 | 85.7 | 3 |
| Q03173 | Protein enabled homolog OS=Mus musculus OX=10090 GN=Enah PE=1 SV=2 | 85.8 | 14 |
| Q99LI8 | Hepatocyte growth factor-regulated tyrosine kinase substrate OS=Mus musculus OX=10090 GN=Hgs PE=1 SV=2 | 86.0 | 3 |
| Q6NZF1 | Zinc finger CCCH domain-containing protein 11A OS=Mus musculus OX=10090 GN=Zc3h11a PE=1 SV=1 | 86.4 | 2 |
| Q924H2 | Mediator of RNA polymerase II transcription subunit 15 OS=Mus musculus OX=10090 GN=Med15 PE=1 SV=3 | 86.6 | 3 |
| Q922K7 | 28S rRNA (cytosine-C(5))-methyltransferase OS=Mus musculus OX=10090 GN=Nop2 PE=1 SV=1 | 86.7 | 2 |
| Q9D4H8 | Cullin-2 OS=Mus musculus OX=10090 GN=Cul2 PE=1 SV=2 | 86.8 | 4 |
| P27612 | Phospholipase A-2-activating protein OS=Mus musculus OX=10090 GN=Plaa PE=1 SV=4 | 87.2 | 2 |
| Q8BML9 | Glutamine--tRNA ligase OS=Mus musculus OX=10090 GN=Qars1 PE=1 SV=1 | 87.6 | 3 |

| | | | |
|---|---|---|---|
| Q8K4Z5 | Splicing factor 3A subunit 1 OS=Mus musculus OX=10090 GN=Sf3a1 PE=1 SV=1 | 88.5 | 14 |
| P25976 | Nucleolar transcription factor 1 OS=Mus musculus OX=10090 GN=Ubtf PE=1 SV=1 | 89.5 | 2 |
| Q62179 | Semaphorin-4B OS=Mus musculus OX=10090 GN=Sema4b PE=1 SV=2 | 91.3 | 9 |
| Q00547 | Hyaluronan mediated motility receptor OS=Mus musculus OX=10090 GN=Hmmr PE=1 SV=4 | 91.7 | 2 |
| Q9WUH7 | Semaphorin-4G OS=Mus musculus OX=10090 GN=Sema4g PE=1 SV=1 | 92.3 | 5 |
| P08113 | Endoplasmin OS=Mus musculus OX=10090 GN=Hsp90b1 PE=1 SV=2 | 92.4 | 2 |
| A2AJI0 | MAP7 domain-containing protein 1 OS=Mus musculus OX=10090 GN=Map7d1 PE=1 SV=1 | 93.2 | 2 |
| Q61316 | Heat shock 70 kDa protein 4 OS=Mus musculus OX=10090 GN=Hspa4 PE=1 SV=1 | 94.1 | 2 |
| Q9Z1X4 | Interleukin enhancer-binding factor 3 OS=Mus musculus OX=10090 GN=Ilf3 PE=1 SV=2 | 96.0 | 3 |
| Q62165 | Dystroglycan 1 OS=Mus musculus OX=10090 GN=Dag1 PE=1 SV=4 | 96.8 | 10 |
| Q3V1V3 | ESF1 homolog OS=Mus musculus OX=10090 GN=Esf1 PE=1 SV=1 | 98.0 | 2 |
| P42567 | Epidermal growth factor receptor substrate 15 OS=Mus musculus OX=10090 GN=Eps15 PE=1 SV=1 | 98.4 | 2 |
| Q9R1E6 | Autotaxin OS=Mus musculus OX=10090 GN=Enpp2 PE=1 SV=3 | 98.8 | 3 |
| Q60902 | Epidermal growth factor receptor substrate 15-like 1 OS=Mus musculus OX=10090 GN=Eps15l1 PE=1 SV=3 | 99.2 | 6 |
| B2RY56 | RNA-binding protein 25 OS=Mus musculus OX=10090 GN=Rbm25 PE=1 SV=2 | 99.5 | 2 |
| P56960 | Exosome complex component 10 OS=Mus musculus OX=10090 GN=Exosc10 PE=1 SV=2 | 100.9 | 2 |
| Q9WU62 | Inner centromere protein OS=Mus musculus OX=10090 GN=Incenp PE=1 SV=2 | 101.1 | 4 |
| Q68FL6 | Methionine--tRNA ligase, cytoplasmic OS=Mus musculus OX=10090 GN=Mars1 PE=1 SV=1 | 101.4 | 5 |
| D3YXK2 | Scaffold attachment factor B1 OS=Mus musculus OX=10090 GN=Safb PE=1 SV=2 | 105.0 | 2 |
| Q0VBL3 | RNA-binding protein 15 OS=Mus musculus OX=10090 GN=Rbm15 PE=1 SV=1 | 105.7 | 5 |
| Q8K019 | Bcl-2-associated transcription factor 1 OS=Mus musculus OX=10090 GN=Bclaf1 PE=1 SV=2 | 105.9 | 7 |
| Q52KI8 | Serine/arginine repetitive matrix protein 1 OS=Mus musculus OX=10090 GN=Srrm1 PE=1 SV=2 | 106.8 | 3 |
| Q569Z6 | Thyroid hormone receptor-associated protein 3 OS=Mus musculus OX=10090 GN=Thrap3 PE=1 SV=1 | 108.1 | 11 |
| Q3UMY5 | Echinoderm microtubule-associated protein-like 4 OS=Mus musculus OX=10090 GN=Eml4 PE=1 SV=1 | 110.0 | 4 |
| A2A432 | Cullin-4B OS=Mus musculus OX=10090 GN=Cul4b PE=1 SV=1 | 110.6 | 3 |
| Q7TQH0 | Ataxin-2-like protein OS=Mus musculus OX=10090 GN=Atxn2l PE=1 SV=1 | 110.6 | 8 |
| O55098 | Serine/threonine-protein kinase 10 OS=Mus musculus OX=10090 GN=Stk10 PE=1 SV=2 | 111.8 | 2 |

| | | | |
|---|---|---|---|
| P11103 | Poly [ADP-ribose] polymerase 1 OS=Mus musculus OX=10090 GN=Parp1 PE=1 SV=3 | 113.0 | 4 |
| O35464 | Semaphorin-6A OS=Mus musculus OX=10090 GN=Sema6a PE=1 SV=2 | 114.4 | 5 |
| O09159 | Lysosomal alpha-mannosidase OS=Mus musculus OX=10090 GN=Man2b1 PE=1 SV=4 | 114.6 | 11 |
| Q9DBR7 | Protein phosphatase 1 regulatory subunit 12A OS=Mus musculus OX=10090 GN=Ppp1r12a PE=1 SV=2 | 114.9 | 2 |
| Q80X50 | Ubiquitin-associated protein 2-like OS=Mus musculus OX=10090 GN=Ubap2l PE=1 SV=1 | 116.7 | 3 |
| P27546 | Microtubule-associated protein 4 OS=Mus musculus OX=10090 GN=Map4 PE=1 SV=3 | 117.4 | 2 |
| Q569Z5 | Probable ATP-dependent RNA helicase DDX46 OS=Mus musculus OX=10090 GN=Ddx46 PE=1 SV=2 | 117.4 | 4 |
| Q91VX2 | Ubiquitin-associated protein 2 OS=Mus musculus OX=10090 GN=Ubap2 PE=1 SV=1 | 117.9 | 5 |
| P34152 | Focal adhesion kinase 1 OS=Mus musculus OX=10090 GN=Ptk2 PE=1 SV=4 | 119.2 | 10 |
| Q920B9 | FACT complex subunit SPT16 OS=Mus musculus OX=10090 GN=Supt16h PE=1 SV=2 | 119.7 | 3 |
| O55201 | Transcription elongation factor SPT5 OS=Mus musculus OX=10090 GN=Supt5h PE=1 SV=1 | 120.6 | 2 |
| Q61147 | Ceruloplasmin OS=Mus musculus OX=10090 GN=Cp PE=1 SV=2 | 121.1 | 4 |
| P42703 | Leukemia inhibitory factor receptor OS=Mus musculus OX=10090 GN=Lifr PE=1 SV=1 | 122.5 | 5 |
| Q80U87 | Ubiquitin carboxyl-terminal hydrolase 8 OS=Mus musculus OX=10090 GN=Usp8 PE=1 SV=2 | 122.5 | 7 |
| Q8CGF7 | Transcription elongation regulator 1 OS=Mus musculus OX=10090 GN=Tcerg1 PE=1 SV=2 | 123.7 | 3 |
| O08784 | Treacle protein OS=Mus musculus OX=10090 GN=Tcof1 PE=1 SV=1 | 134.9 | 3 |
| O70305 | Ataxin-2 OS=Mus musculus OX=10090 GN=Atxn2 PE=1 SV=1 | 136.4 | 4 |
| Q05D44 | Eukaryotic translation initiation factor 5B OS=Mus musculus OX=10090 GN=Eif5b PE=1 SV=2 | 137.5 | 2 |
| Q9EPX2 | Papilin OS=Mus musculus OX=10090 GN=Papln PE=2 SV=2 | 138.8 | 19 |
| P33174 | Chromosome-associated kinesin KIF4 OS=Mus musculus OX=10090 GN=Kif4 PE=1 SV=3 | 139.4 | 3 |
| Q6ZPZ3 | Zinc finger CCCH domain-containing protein 4 OS=Mus musculus OX=10090 GN=Zc3h4 PE=1 SV=2 | 140.9 | 8 |
| Q62417 | Sorbin and SH3 domain-containing protein 1 OS=Mus musculus OX=10090 GN=Sorbs1 PE=1 SV=2 | 143.0 | 5 |
| Q99KY4 | Cyclin-G-associated kinase OS=Mus musculus OX=10090 GN=Gak PE=1 SV=2 | 143.6 | 3 |
| Q8K4P0 | pre-mRNA 3' end processing protein WDR33 OS=Mus musculus OX=10090 GN=Wdr33 PE=1 SV=1 | 145.2 | 5 |
| Q8CG47 | Structural maintenance of chromosomes protein 4 OS=Mus musculus OX=10090 GN=Smc4 PE=1 SV=1 | 146.8 | 4 |
| Q9JIX8 | Apoptotic chromatin condensation inducer in the nucleus OS=Mus musculus OX=10090 GN=Acin1 PE=1 SV=3 | 150.6 | 4 |
| Q7TPV4 | Myb-binding protein 1A OS=Mus musculus OX=10090 GN=Mybbp1a PE=1 SV=2 | 151.9 | 2 |
| Q61595 | Kinectin OS=Mus musculus OX=10090 GN=Ktn1 PE=1 SV=1 | 152.5 | 2 |

| | | | |
|---|---|---|---|
| Q5FWI3 | Cell surface hyaluronidase OS=Mus musculus OX=10090 GN=Cemip2 PE=1 SV=1 | 153.7 | 2 |
| Q922J3 | CAP-Gly domain-containing linker protein 1 OS=Mus musculus OX=10090 GN=Clip1 PE=1 SV=1 | 155.7 | 3 |
| Q9ESU6 | Bromodomain-containing protein 4 OS=Mus musculus OX=10090 GN=Brd4 PE=1 SV=2 | 155.8 | 2 |
| Q9EQJ9 | Membrane-associated guanylate kinase, WW and PDZ domain-containing protein 3 OS=Mus musculus OX=10090 GN=Magi3 PE=1 SV=2 | 161.6 | 2 |
| P23116 | Eukaryotic translation initiation factor 3 subunit A OS=Mus musculus OX=10090 GN=Eif3a PE=1 SV=5 | 161.8 | 6 |
| Q8CGC7 | Bifunctional glutamate/proline--tRNA ligase OS=Mus musculus OX=10090 GN=Eprs1 PE=1 SV=4 | 170.0 | 9 |
| Q99PL5 | Ribosome-binding protein 1 OS=Mus musculus OX=10090 GN=Rrbp1 PE=1 SV=2 | 172.8 | 8 |
| P13864 | DNA (cytosine-5)-methyltransferase 1 OS=Mus musculus OX=10090 GN=Dnmt1 PE=1 SV=5 | 183.1 | 4 |
| B2RX14 | Terminal uridylyltransferase 4 OS=Mus musculus OX=10090 GN=Tut4 PE=1 SV=2 | 184.5 | 4 |
| Q9JKF1 | Ras GTPase-activating-like protein IQGAP1 OS=Mus musculus OX=10090 GN=Iqgap1 PE=1 SV=2 | 188.6 | 4 |
| Q9Z0R4 | Intersectin-1 OS=Mus musculus OX=10090 GN=Itsn1 PE=1 SV=2 | 194.2 | 4 |
| P97868 | E3 ubiquitin-protein ligase RBBP6 OS=Mus musculus OX=10090 GN=Rbbp6 PE=1 SV=5 | 199.5 | 2 |
| A2ASQ1 | Agrin OS=Mus musculus OX=10090 GN=Agrn PE=1 SV=1 | 207.4 | 2 |
| B0V2N1 | Receptor-type tyrosine-protein phosphatase S OS=Mus musculus OX=10090 GN=Ptprs PE=1 SV=1 | 211.8 | 5 |
| Q80UG2 | Plexin-A4 OS=Mus musculus OX=10090 GN=Plxna4 PE=1 SV=3 | 212.4 | 6 |
| Q8VDD5 | Myosin-9 OS=Mus musculus OX=10090 GN=Myh9 PE=1 SV=4 | 226.2 | 3 |
| B2RWS6 | Histone acetyltransferase p300 OS=Mus musculus OX=10090 GN=Ep300 PE=1 SV=2 | 263.1 | 2 |
| F6ZDS4 | Nucleoprotein TPR OS=Mus musculus OX=10090 GN=Tpr PE=1 SV=1 | 273.8 | 11 |
| Q62261 | Spectrin beta chain, non-erythrocytic 1 OS=Mus musculus OX=10090 GN=Sptbn1 PE=1 SV=2 | 274.1 | 2 |
| Q8BTM8 | Filamin-A OS=Mus musculus OX=10090 GN=Flna PE=1 SV=5 | 281.0 | 15 |
| Q811L6 | Microtubule-associated serine/threonine-protein kinase 4 OS=Mus musculus OX=10090 GN=Mast4 PE=1 SV=3 | 283.8 | 6 |
| Q8BTI8 | Serine/arginine repetitive matrix protein 2 OS=Mus musculus OX=10090 GN=Srrm2 PE=1 SV=3 | 294.7 | 3 |
| Q9QYR6 | Microtubule-associated protein 1A OS=Mus musculus OX=10090 GN=Map1a PE=1 SV=2 | 300.0 | 2 |
| Q3TLH4 | Protein PRRC2C OS=Mus musculus OX=10090 GN=Prrc2c PE=1 SV=3 | 310.7 | 13 |
| Q6KCD5 | Nipped-B-like protein OS=Mus musculus OX=10090 GN=Nipbl PE=1 SV=1 | 315.3 | 8 |
| E9Q6J5 | Biorientation of chromosomes in cell division protein 1-like 1 OS=Mus musculus OX=10090 GN=Bod1l PE=1 SV=1 | 327.3 | 2 |
| E9PVX6 | Proliferation marker protein Ki-67 OS=Mus musculus OX=10090 GN=Mki67 PE=1 SV=1 | 350.7 | 2 |
| Q62504 | Msx2-interacting protein OS=Mus musculus OX=10090 GN=Spen PE=1 SV=2 | 398.5 | 4 |

| | | | |
|---|---|---|---|
| Q8R0W0 | Epiplakin OS=Mus musculus OX=10090 GN=Eppk1 PE=1 SV=2 | 724.2 | 2 |

(**B**)

| Accession | Description | MW, kDa | # Unique Peptides |
|---|---|---|---|
| P62858 | Small ribosomal subunit protein eS28 OS=Mus musculus OX=10090 GN=Rps28 PE=1 SV=1 | 7.8 | 3 |
| Q9JIX0 | Transcription and mRNA export factor ENY2 OS=Mus musculus OX=10090 GN=Eny2 PE=1 SV=1 | 11.5 | 2 |
| P62897 | Cytochrome c, somatic OS=Mus musculus OX=10090 GN=Cycs PE=1 SV=2 | 11.6 | 3 |
| O35143 | ATPase inhibitor, mitochondrial OS=Mus musculus OX=10090 GN=Atp5if1 PE=1 SV=2 | 12.2 | 4 |
| P83870 | PHD finger-like domain-containing protein 5A OS=Mus musculus OX=10090 GN=Phf5a PE=1 SV=1 | 12.4 | 3 |
| P70349 | Adenosine 5'-monophosphoramidase HINT1 OS=Mus musculus OX=10090 GN=Hint1 PE=1 SV=3 | 13.8 | 2 |
| P01887 | Beta-2-microglobulin OS=Mus musculus OX=10090 GN=B2m PE=1 SV=2 | 13.8 | 3 |
| P56812 | Programmed cell death protein 5 OS=Mus musculus OX=10090 GN=Pdcd5 PE=1 SV=3 | 14.3 | 6 |
| P63323 | Small ribosomal subunit protein eS12 OS=Mus musculus OX=10090 GN=Rps12 PE=1 SV=3 | 14.5 | 2 |
| P45878 | Peptidyl-prolyl cis-trans isomerase FKBP2 OS=Mus musculus OX=10090 GN=Fkbp2 PE=1 SV=1 | 15.3 | 7 |
| P21460 | Cystatin-C OS=Mus musculus OX=10090 GN=Cst3 PE=1 SV=2 | 15.5 | 4 |
| Q8VHC3 | Selenoprotein M OS=Mus musculus OX=10090 GN=Selenom PE=1 SV=3 | 16.4 | 4 |
| Q9D2M8 | Ubiquitin-conjugating enzyme E2 variant 2 OS=Mus musculus OX=10090 GN=Ube2v2 PE=1 SV=4 | 16.4 | 4 |
| Q8R3Q6 | Protein MIX23 OS=Mus musculus OX=10090 GN=Mix23 PE=1 SV=1 | 16.7 | 3 |
| P62838 | Ubiquitin-conjugating enzyme E2 D2 OS=Mus musculus OX=10090 GN=Ube2d2 PE=1 SV=1 | 16.7 | 2 |
| O70200 | Allograft inflammatory factor 1 OS=Mus musculus OX=10090 GN=Aif1 PE=1 SV=1 | 16.9 | 2 |
| P54227 | Stathmin OS=Mus musculus OX=10090 GN=Stmn1 PE=1 SV=2 | 17.3 | 6 |
| Q01768 | Nucleoside diphosphate kinase B OS=Mus musculus OX=10090 GN=Nme2 PE=1 SV=1 | 17.4 | 7 |
| P62983 | Ubiquitin-ribosomal protein eS31 fusion protein OS=Mus musculus OX=10090 GN=Rps27a PE=1 SV=2 | 17.9 | 3 |
| P17742 | Peptidyl-prolyl cis-trans isomerase A OS=Mus musculus OX=10090 GN=Ppia PE=1 SV=2 | 18.0 | 3 |
| Q9D7P6 | Iron-sulfur cluster assembly enzyme ISCU OS=Mus musculus OX=10090 GN=Iscu PE=1 SV=1 | 18.1 | 5 |
| P18760 | Cofilin-1 OS=Mus musculus OX=10090 GN=Cfl1 PE=1 SV=3 | 18.5 | 3 |
| Q8R2Q8 | Bone marrow stromal antigen 2 OS=Mus musculus OX=10090 GN=Bst2 PE=1 SV=1 | 19.1 | 2 |
| P53996 | CCHC-type zinc finger nucleic acid binding protein OS=Mus musculus OX=10090 GN=Cnbp PE=1 SV=2 | 19.6 | 3 |
| Q9D1M4 | Eukaryotic translation elongation factor 1 epsilon-1 OS=Mus musculus OX=10090 GN=Eef1e1 PE=1 SV=1 | 19.8 | 5 |
| Q9DB15 | Large ribosomal subunit protein bL12m OS=Mus musculus OX=10090 GN=Mrpl12 PE=1 SV=2 | 21.7 | 4 |

| | | | |
|---|---|---|---|
| Q99KR7 | Peptidyl-prolyl cis-trans isomerase F, mitochondrial OS=Mus musculus OX=10090 GN=Ppif PE=1 SV=1 | 21.7 | 3 |
| O88952 | Protein lin-7 homolog C OS=Mus musculus OX=10090 GN=Lin7c PE=1 SV=2 | 21.8 | 2 |
| P99029 | Peroxiredoxin-5, mitochondrial OS=Mus musculus OX=10090 GN=Prdx5 PE=1 SV=2 | 21.9 | 6 |
| Q9DBP5 | UMP-CMP kinase OS=Mus musculus OX=10090 GN=Cmpk1 PE=1 SV=1 | 22.2 | 6 |
| P35700 | Peroxiredoxin-1 OS=Mus musculus OX=10090 GN=Prdx1 PE=1 SV=1 | 22.2 | 7 |
| Q923D2 | Flavin reductase (NADPH) OS=Mus musculus OX=10090 GN=Blvrb PE=1 SV=3 | 22.2 | 7 |
| Q9WVA4 | Transgelin-2 OS=Mus musculus OX=10090 GN=Tagln2 PE=1 SV=4 | 22.4 | 9 |
| P56213 | FAD-linked sulfhydryl oxidase ALR OS=Mus musculus OX=10090 GN=Gfer PE=1 SV=2 | 22.9 | 4 |
| Q9DCT5 | Stromal cell-derived factor 2 OS=Mus musculus OX=10090 GN=Sdf2 PE=1 SV=1 | 23.1 | 2 |
| Q6IRU5-2 | Isoform 2 of Clathrin light chain B OS=Mus musculus OX=10090 GN=Cltb | 23.2 | 5 |
| P19157 | Glutathione S-transferase P 1 OS=Mus musculus OX=10090 GN=Gstp1 PE=1 SV=2 | 23.6 | 4 |
| P24369 | Peptidyl-prolyl cis-trans isomerase B OS=Mus musculus OX=10090 GN=Ppib PE=1 SV=2 | 23.7 | 3 |
| P30681 | High mobility group protein B2 OS=Mus musculus OX=10090 GN=Hmgb2 PE=1 SV=3 | 24.1 | 5 |
| O35166 | Golgi SNAP receptor complex member 2 OS=Mus musculus OX=10090 GN=Gosr2 PE=1 SV=2 | 24.7 | 5 |
| O08709 | Peroxiredoxin-6 OS=Mus musculus OX=10090 GN=Prdx6 PE=1 SV=3 | 24.9 | 3 |
| Q9D8B3 | Charged multivesicular body protein 4b OS=Mus musculus OX=10090 GN=Chmp4b PE=1 SV=2 | 24.9 | 6 |
| Q62446 | Peptidyl-prolyl cis-trans isomerase FKBP3 OS=Mus musculus OX=10090 GN=Fkbp3 PE=1 SV=2 | 25.1 | 2 |
| Q60631 | Growth factor receptor-bound protein 2 OS=Mus musculus OX=10090 GN=Grb2 PE=1 SV=1 | 25.2 | 2 |
| Q9CQI7 | U2 small nuclear ribonucleoprotein B'' OS=Mus musculus OX=10090 GN=Snrpb2 PE=1 SV=1 | 25.3 | 2 |
| Q62093 | Serine/arginine-rich splicing factor 2 OS=Mus musculus OX=10090 GN=Srsf2 PE=1 SV=4 | 25.5 | 2 |
| Q61206 | Platelet-activating factor acetylhydrolase IB subunit alpha2 OS=Mus musculus OX=10090 GN=Pafah1b2 PE=1 SV=2 | 25.6 | 2 |
| Q61205 | Platelet-activating factor acetylhydrolase IB subunit alpha1 OS=Mus musculus OX=10090 GN=Pafah1b3 PE=1 SV=1 | 25.8 | 5 |
| Q9CQF3 | Cleavage and polyadenylation specificity factor subunit 5 OS=Mus musculus OX=10090 GN=Nudt21 PE=1 SV=1 | 26.2 | 6 |
| Q9CXW3 | Calcyclin-binding protein OS=Mus musculus OX=10090 GN=Cacybp PE=1 SV=1 | 26.5 | 2 |
| Q9WTP6 | Adenylate kinase 2, mitochondrial OS=Mus musculus OX=10090 GN=Ak2 PE=1 SV=5 | 26.5 | 9 |
| O08583 | THO complex subunit 4 OS=Mus musculus OX=10090 GN=Alyref PE=1 SV=3 | 26.9 | 3 |
| Q03402 | Cysteine-rich secretory protein 3 OS=Mus musculus OX=10090 GN=Crisp3 PE=1 SV=1 | 27.3 | 5 |

| | | | |
|---|---|---|---|
| Q6PDM2 | Serine/arginine-rich splicing factor 1 OS=Mus musculus OX=10090 GN=Srsf1 PE=1 SV=3 | 27.7 | 4 |
| Q9WV55 | Vesicle-associated membrane protein-associated protein A OS=Mus musculus OX=10090 GN=Vapa PE=1 SV=2 | 27.8 | 4 |
| Q61335 | B-cell receptor-associated protein 31 OS=Mus musculus OX=10090 GN=Bcap31 PE=1 SV=4 | 27.9 | 2 |
| Q9CQE1 | Protein NipSnap homolog 3B OS=Mus musculus OX=10090 GN=Nipsnap3b PE=1 SV=1 | 28.3 | 6 |
| Q9DBJ1 | Phosphoglycerate mutase 1 OS=Mus musculus OX=10090 GN=Pgam1 PE=1 SV=3 | 28.8 | 2 |
| Q99020 | Heterogeneous nuclear ribonucleoprotein A/B OS=Mus musculus OX=10090 GN=Hnrnpab PE=1 SV=1 | 30.8 | 3 |
| Q8BL97 | Serine/arginine-rich splicing factor 7 OS=Mus musculus OX=10090 GN=Srsf7 PE=1 SV=1 | 30.8 | 6 |
| O35326 | Serine/arginine-rich splicing factor 5 OS=Mus musculus OX=10090 GN=Srsf5 PE=1 SV=2 | 30.9 | 2 |
| Q9ER00 | Syntaxin-12 OS=Mus musculus OX=10090 GN=Stx12 PE=1 SV=1 | 31.2 | 8 |
| Q9R0Q4 | Mortality factor 4-like protein 2 OS=Mus musculus OX=10090 GN=Morf4l2 PE=1 SV=1 | 32.2 | 5 |
| Q61937 | Nucleophosmin OS=Mus musculus OX=10090 GN=Npm1 PE=1 SV=1 | 32.5 | 4 |
| Q8K4F5 | sn-1-specific diacylglycerol lipase ABHD11 OS=Mus musculus OX=10090 GN=Abhd11 PE=1 SV=1 | 33.5 | 9 |
| Q8C0M9 | Isoaspartyl peptidase/L-asparaginase OS=Mus musculus OX=10090 GN=Asrgl1 PE=1 SV=1 | 33.9 | 3 |
| Q9WUU7 | Cathepsin Z OS=Mus musculus OX=10090 GN=Ctsz PE=1 SV=1 | 34.0 | 2 |
| P31230 | Aminoacyl tRNA synthase complex-interacting multifunctional protein 1 OS=Mus musculus OX=10090 GN=Aimp1 PE=1 SV=2 | 34.0 | 2 |
| P49312 | Heterogeneous nuclear ribonucleoprotein A1 OS=Mus musculus OX=10090 GN=Hnrnpa1 PE=1 SV=2 | 34.2 | 7 |
| O88531 | Palmitoyl-protein thioesterase 1 OS=Mus musculus OX=10090 GN=Ppt1 PE=1 SV=2 | 34.5 | 6 |
| Q91YR9 | Prostaglandin reductase 1 OS=Mus musculus OX=10090 GN=Ptgr1 PE=1 SV=2 | 35.5 | 10 |
| Q9Z1D1 | Eukaryotic translation initiation factor 3 subunit G OS=Mus musculus OX=10090 GN=Eif3g PE=1 SV=2 | 35.6 | 2 |
| P08249 | Malate dehydrogenase, mitochondrial OS=Mus musculus OX=10090 GN=Mdh2 PE=1 SV=3 | 35.6 | 10 |
| P70372 | ELAV-like protein 1 OS=Mus musculus OX=10090 GN=Elavl1 PE=1 SV=2 | 36.1 | 6 |
| P06151 | L-lactate dehydrogenase A chain OS=Mus musculus OX=10090 GN=Ldha PE=1 SV=3 | 36.5 | 3 |
| P08101 | Low affinity immunoglobulin gamma Fc region receptor II OS=Mus musculus OX=10090 GN=Fcgr2 PE=1 SV=2 | 36.7 | 5 |
| Q93092 | Transaldolase OS=Mus musculus OX=10090 GN=Taldo1 PE=1 SV=2 | 37.4 | 10 |
| O88569 | Heterogeneous nuclear ribonucleoproteins A2/B1 OS=Mus musculus OX=10090 GN=Hnrnpa2b1 PE=1 SV=2 | 37.4 | 7 |
| P06797 | Procathepsin L OS=Mus musculus OX=10090 GN=Ctsl PE=1 SV=2 | 37.5 | 2 |
| P60335 | Poly(rC)-binding protein 1 OS=Mus musculus OX=10090 GN=Pcbp1 PE=1 SV=1 | 37.5 | 5 |

| | | | |
|---|---|---|---|
| P70441 | Na(+)/H(+) exchange regulatory cofactor NHE-RF1 OS=Mus musculus OX=10090 GN=Nherf1 PE=1 SV=3 | 38.6 | 5 |
| P56542 | Deoxyribonuclease-2-alpha OS=Mus musculus OX=10090 GN=Dnase2 PE=1 SV=1 | 38.8 | 6 |
| Q9JKB3 | Y-box-binding protein 3 OS=Mus musculus OX=10090 GN=Ybx3 PE=1 SV=2 | 38.8 | 2 |
| P05064 | Fructose-bisphosphate aldolase A OS=Mus musculus OX=10090 GN=Aldoa PE=1 SV=2 | 39.3 | 31 |
| P05063 | Fructose-bisphosphate aldolase C OS=Mus musculus OX=10090 GN=Aldoc PE=1 SV=4 | 39.4 | 24 |
| P28474 | Alcohol dehydrogenase class-3 OS=Mus musculus OX=10090 GN=Adh5 PE=1 SV=3 | 39.5 | 2 |
| P59481 | VIP36-like protein OS=Mus musculus OX=10090 GN=Lman2l PE=1 SV=1 | 39.9 | 5 |
| Q91YR1 | Twinfilin-1 OS=Mus musculus OX=10090 GN=Twf1 PE=1 SV=2 | 40.1 | 4 |
| Q78JW9 | Ubiquitin domain-containing protein UBFD1 OS=Mus musculus OX=10090 GN=Ubfd1 PE=1 SV=2 | 40.1 | 2 |
| Q9CZ44 | NSFL1 cofactor p47 OS=Mus musculus OX=10090 GN=Nsfl1c PE=1 SV=1 | 40.7 | 13 |
| Q9CR16 | Peptidyl-prolyl cis-trans isomerase D OS=Mus musculus OX=10090 GN=Ppid PE=1 SV=3 | 40.7 | 6 |
| Q91VM5 | RNA binding motif protein, X-linked-like-1 OS=Mus musculus OX=10090 GN=Rbmxl1 PE=2 SV=1 | 42.1 | 4 |
| P34902 | Cytokine receptor common subunit gamma OS=Mus musculus OX=10090 GN=Il2rg PE=1 SV=1 | 42.2 | 4 |
| P55302 | Alpha-2-macroglobulin receptor-associated protein OS=Mus musculus OX=10090 GN=Lrpap1 PE=1 SV=1 | 42.2 | 6 |
| P70318 | Nucleolysin TIAR OS=Mus musculus OX=10090 GN=Tial1 PE=1 SV=1 | 43.4 | 4 |
| P04202 | Transforming growth factor beta-1 proprotein OS=Mus musculus OX=10090 GN=Tgfb1 PE=1 SV=1 | 44.3 | 3 |
| P15535 | Beta-1,4-galactosyltransferase 1 OS=Mus musculus OX=10090 GN=B4galt1 PE=1 SV=1 | 44.4 | 2 |
| P09411 | Phosphoglycerate kinase 1 OS=Mus musculus OX=10090 GN=Pgk1 PE=1 SV=4 | 44.5 | 8 |
| Q9CY58 | SERPINE1 mRNA-binding protein 1 OS=Mus musculus OX=10090 GN=Serbp1 PE=1 SV=2 | 44.7 | 2 |
| P18242 | Cathepsin D OS=Mus musculus OX=10090 GN=Ctsd PE=1 SV=1 | 44.9 | 8 |
| Q64685 | Beta-galactoside alpha-2,6-sialyltransferase 1 OS=Mus musculus OX=10090 GN=St6gal1 PE=1 SV=2 | 46.6 | 2 |
| P17182 | Alpha-enolase OS=Mus musculus OX=10090 GN=Eno1 PE=1 SV=3 | 47.1 | 3 |
| Q922R8 | Protein disulfide-isomerase A6 OS=Mus musculus OX=10090 GN=Pdia6 PE=1 SV=3 | 48.1 | 12 |
| Q62418 | Drebrin-like protein OS=Mus musculus OX=10090 GN=Dbnl PE=1 SV=2 | 48.7 | 6 |
| O35737 | Heterogeneous nuclear ribonucleoprotein H OS=Mus musculus OX=10090 GN=Hnrnph1 PE=1 SV=3 | 49.2 | 4 |
| P11680 | Properdin OS=Mus musculus OX=10090 GN=Cfp PE=1 SV=2 | 50.3 | 6 |
| O55131 | Septin-7 OS=Mus musculus OX=10090 GN=Septin7 PE=1 SV=1 | 50.5 | 4 |

| | | | |
|---|---|---|---|
| Q80V42 | Carboxypeptidase M OS=Mus musculus OX=10090 GN=Cpm PE=1 SV=2 | 50.5 | 2 |
| Q9QUN3 | B-cell linker protein OS=Mus musculus OX=10090 GN=Blnk PE=1 SV=1 | 50.6 | 4 |
| P40124 | Adenylyl cyclase-associated protein 1 OS=Mus musculus OX=10090 GN=Cap1 PE=1 SV=4 | 51.5 | 5 |
| Q64287 | Interferon regulatory factor 4 OS=Mus musculus OX=10090 GN=Irf4 PE=1 SV=1 | 51.5 | 9 |
| P97855 | Ras GTPase-activating protein-binding protein 1 OS=Mus musculus OX=10090 GN=G3bp1 PE=1 SV=1 | 51.8 | 3 |
| Q6NXH2 | Glycoprotein endo-alpha-1,2-mannosidase OS=Mus musculus OX=10090 GN=Manea PE=2 SV=1 | 53.1 | 6 |
| Q02819 | Nucleobindin-1 OS=Mus musculus OX=10090 GN=Nucb1 PE=1 SV=2 | 53.4 | 3 |
| P17892 | Pancreatic lipase-related protein 2 OS=Mus musculus OX=10090 GN=Pnliprp2 PE=1 SV=2 | 54.0 | 10 |
| P49710 | Hematopoietic lineage cell-specific protein OS=Mus musculus OX=10090 GN=Hcls1 PE=1 SV=2 | 54.2 | 4 |
| P97807 | Fumarate hydratase, mitochondrial OS=Mus musculus OX=10090 GN=Fh PE=1 SV=3 | 54.3 | 8 |
| Q99K48 | Non-POU domain-containing octamer-binding protein OS=Mus musculus OX=10090 GN=Nono PE=1 SV=3 | 54.5 | 3 |
| Q02853 | Stromelysin-3 OS=Mus musculus OX=10090 GN=Mmp11 PE=1 SV=2 | 55.4 | 2 |
| Q9QYI3 | DnaJ homolog subfamily C member 7 OS=Mus musculus OX=10090 GN=Dnajc7 PE=1 SV=2 | 56.4 | 4 |
| Q8VCH8 | UBX domain-containing protein 4 OS=Mus musculus OX=10090 GN=Ubxn4 PE=1 SV=1 | 56.4 | 4 |
| P97360 | Transcription factor ETV6 OS=Mus musculus OX=10090 GN=Etv6 PE=1 SV=1 | 56.4 | 3 |
| O35664 | Interferon alpha/beta receptor 2 OS=Mus musculus OX=10090 GN=Ifnar2 PE=1 SV=2 | 56.5 | 2 |
| P09242 | Alkaline phosphatase, tissue-nonspecific isozyme OS=Mus musculus OX=10090 GN=Alpl PE=1 SV=2 | 57.5 | 3 |
| Q3UEB3-2 | Isoform 2 of Poly(U)-binding-splicing factor PUF60 OS=Mus musculus OX=10090 GN=Puf60 | 58.5 | 6 |
| Q6NVF9 | Cleavage and polyadenylation specificity factor subunit 6 OS=Mus musculus OX=10090 GN=Cpsf6 PE=1 SV=1 | 59.1 | 4 |
| Q9Z2A5 | Arginyl-tRNA--protein transferase 1 OS=Mus musculus OX=10090 GN=Ate1 PE=1 SV=2 | 59.1 | 3 |
| P32020 | Sterol carrier protein 2 OS=Mus musculus OX=10090 GN=Scp2 PE=1 SV=3 | 59.1 | 10 |
| P17225 | Polypyrimidine tract-binding protein 1 OS=Mus musculus OX=10090 GN=Ptbp1 PE=1 SV=3 | 59.3 | 3 |
| Q99JF8 | PC4 and SFRS1-interacting protein OS=Mus musculus OX=10090 GN=Psip1 PE=1 SV=1 | 59.7 | 2 |
| Q8CI11 | Guanine nucleotide-binding protein-like 3 OS=Mus musculus OX=10090 GN=Gnl3 PE=1 SV=2 | 60.7 | 3 |
| P20060 | Beta-hexosaminidase subunit beta OS=Mus musculus OX=10090 GN=Hexb PE=1 SV=2 | 61.1 | 8 |
| Q60864 | Stress-induced-phosphoprotein 1 OS=Mus musculus OX=10090 GN=Stip1 PE=1 SV=1 | 62.5 | 12 |
| P03975 | IgE-binding protein OS=Mus musculus OX=10090 GN=Iap PE=2 SV=1 | 62.7 | 6 |

| | | | |
|---|---|---|---|
| P06745 | Glucose-6-phosphate isomerase OS=Mus musculus OX=10090 GN=Gpi PE=1 SV=4 | 62.7 | 21 |
| Q9D2L1 | Arylsulfatase K OS=Mus musculus OX=10090 GN=Arsk PE=1 SV=2 | 62.8 | 3 |
| Q8C854 | Myelin expression factor 2 OS=Mus musculus OX=10090 GN=Myef2 PE=1 SV=1 | 63.3 | 3 |
| Q8CHU3 | Epsin-2 OS=Mus musculus OX=10090 GN=Epn2 PE=1 SV=1 | 63.4 | 4 |
| P28798 | Progranulin OS=Mus musculus OX=10090 GN=Grn PE=1 SV=2 | 63.4 | 4 |
| Q5NCR9 | Nuclear speckle splicing regulatory protein 1 OS=Mus musculus OX=10090 GN=Nsrp1 PE=1 SV=1 | 63.8 | 2 |
| Q80WJ7 | Protein LYRIC OS=Mus musculus OX=10090 GN=Mtdh PE=1 SV=1 | 63.8 | 2 |
| Q6PB93 | Polypeptide N-acetylgalactosaminyltransferase 2 OS=Mus musculus OX=10090 GN=Galnt2 PE=1 SV=1 | 64.5 | 18 |
| Q60862 | Origin recognition complex subunit 2 OS=Mus musculus OX=10090 GN=Orc2 PE=1 SV=1 | 65.9 | 2 |
| Q9Z0X1 | Apoptosis-inducing factor 1, mitochondrial OS=Mus musculus OX=10090 GN=Aifm1 PE=1 SV=1 | 66.7 | 13 |
| P40142 | Transketolase OS=Mus musculus OX=10090 GN=Tkt PE=1 SV=1 | 67.6 | 15 |
| Q61545 | RNA-binding protein EWS OS=Mus musculus OX=10090 GN=Ewsr1 PE=1 SV=2 | 68.4 | 3 |
| Q99KN9 | Clathrin interactor 1 OS=Mus musculus OX=10090 GN=Clint1 PE=1 SV=2 | 68.5 | 7 |
| Q8BGD9 | Eukaryotic translation initiation factor 4B OS=Mus musculus OX=10090 GN=Eif4b PE=1 SV=1 | 68.8 | 7 |
| Q9DBG7 | Signal recognition particle receptor subunit alpha OS=Mus musculus OX=10090 GN=Srpra PE=1 SV=1 | 69.6 | 2 |
| Q5RKZ7 | Molybdenum cofactor biosynthesis protein 1 OS=Mus musculus OX=10090 GN=Mocs1 PE=1 SV=2 | 69.8 | 4 |
| Q64213 | Splicing factor 1 OS=Mus musculus OX=10090 GN=Sf1 PE=1 SV=6 | 70.4 | 2 |
| Q9JLQ0 | CD2-associated protein OS=Mus musculus OX=10090 GN=Cd2ap PE=1 SV=3 | 70.4 | 4 |
| P29341 | Polyadenylate-binding protein 1 OS=Mus musculus OX=10090 GN=Pabpc1 PE=1 SV=2 | 70.6 | 10 |
| P63017 | Heat shock cognate 71 kDa protein OS=Mus musculus OX=10090 GN=Hspa8 PE=1 SV=1 | 70.8 | 11 |
| Q8K2Q9 | Shootin-1 OS=Mus musculus OX=10090 GN=Shtn1 PE=1 SV=1 | 71.3 | 2 |
| Q3U9G9 | Delta(14)-sterol reductase LBR OS=Mus musculus OX=10090 GN=Lbr PE=1 SV=2 | 71.4 | 2 |
| Q8C7U7 | Polypeptide N-acetylgalactosaminyltransferase 6 OS=Mus musculus OX=10090 GN=Galnt6 PE=2 SV=1 | 71.5 | 19 |
| P54103 | DnaJ homolog subfamily C member 2 OS=Mus musculus OX=10090 GN=Dnajc2 PE=1 SV=2 | 71.7 | 2 |
| Q9D706 | RNA polymerase II-associated protein 3 OS=Mus musculus OX=10090 GN=Rpap3 PE=1 SV=1 | 74.1 | 6 |
| P48678 | Prelamin-A/C OS=Mus musculus OX=10090 GN=Lmna PE=1 SV=2 | 74.2 | 4 |
| Q9QUR8 | Semaphorin-7A OS=Mus musculus OX=10090 GN=Sema7a PE=1 SV=1 | 74.9 | 13 |

| | | | |
|---|---|---|---|
| Q8VIJ6 | Splicing factor, proline- and glutamine-rich OS=Mus musculus OX=10090 GN=Sfpq PE=1 SV=1 | 75.4 | 2 |
| P09405 | Nucleolin OS=Mus musculus OX=10090 GN=Ncl PE=1 SV=2 | 76.7 | 2 |
| Q8BWW4 | La-related protein 4 OS=Mus musculus OX=10090 GN=Larp4 PE=1 SV=2 | 79.7 | 2 |
| O88967 | ATP-dependent zinc metalloprotease YME1L1 OS=Mus musculus OX=10090 GN=Yme1l1 PE=1 SV=1 | 80.0 | 3 |
| P26928 | Hepatocyte growth factor-like protein OS=Mus musculus OX=10090 GN=Mst1 PE=2 SV=2 | 80.6 | 13 |
| Q6A0A2 | La-related protein 4B OS=Mus musculus OX=10090 GN=Larp4b PE=1 SV=2 | 81.6 | 8 |
| Q8BND5 | Sulfhydryl oxidase 1 OS=Mus musculus OX=10090 GN=Qsox1 PE=1 SV=1 | 82.7 | 11 |
| Q9D0R2 | Threonine--tRNA ligase 1, cytoplasmic OS=Mus musculus OX=10090 GN=Tars1 PE=1 SV=2 | 83.3 | 2 |
| P51125 | Calpastatin OS=Mus musculus OX=10090 GN=Cast PE=1 SV=2 | 84.9 | 4 |
| Q68FF6 | ARF GTPase-activating protein GIT1 OS=Mus musculus OX=10090 GN=Git1 PE=1 SV=1 | 85.2 | 4 |
| Q03173 | Protein enabled homolog OS=Mus musculus OX=10090 GN=Enah PE=1 SV=2 | 85.8 | 8 |
| Q9D4H8 | Cullin-2 OS=Mus musculus OX=10090 GN=Cul2 PE=1 SV=2 | 86.8 | 2 |
| Q8K4Z5 | Splicing factor 3A subunit 1 OS=Mus musculus OX=10090 GN=Sf3a1 PE=1 SV=1 | 88.5 | 9 |
| Q8BLY2 | Threonine--tRNA ligase 2, cytoplasmic OS=Mus musculus OX=10090 GN=Tars3 PE=1 SV=1 | 91.3 | 2 |
| Q62179 | Semaphorin-4B OS=Mus musculus OX=10090 GN=Sema4b PE=1 SV=2 | 91.3 | 10 |
| A2AJI0 | MAP7 domain-containing protein 1 OS=Mus musculus OX=10090 GN=Map7d1 PE=1 SV=1 | 93.2 | 2 |
| Q8C0D4 | Rho GTPase-activating protein 12 OS=Mus musculus OX=10090 GN=Arhgap12 PE=1 SV=2 | 95.3 | 2 |
| O09126 | Semaphorin-4D OS=Mus musculus OX=10090 GN=Sema4d PE=1 SV=2 | 95.6 | 2 |
| Q9Z1X4 | Interleukin enhancer-binding factor 3 OS=Mus musculus OX=10090 GN=Ilf3 PE=1 SV=2 | 96.0 | 4 |
| Q62165 | Dystroglycan 1 OS=Mus musculus OX=10090 GN=Dag1 PE=1 SV=4 | 96.8 | 9 |
| Q3V1V3 | ESF1 homolog OS=Mus musculus OX=10090 GN=Esf1 PE=1 SV=1 | 98.0 | 2 |
| Q60902 | Epidermal growth factor receptor substrate 15-like 1 OS=Mus musculus OX=10090 GN=Eps15l1 PE=1 SV=3 | 99.2 | 3 |
| Q9WU62 | Inner centromere protein OS=Mus musculus OX=10090 GN=Incenp PE=1 SV=2 | 101.1 | 5 |
| Q68FL6 | Methionine--tRNA ligase, cytoplasmic OS=Mus musculus OX=10090 GN=Mars1 PE=1 SV=1 | 101.4 | 3 |
| D3YXK2 | Scaffold attachment factor B1 OS=Mus musculus OX=10090 GN=Safb PE=1 SV=2 | 105.0 | 3 |
| Q0VBL3 | RNA-binding protein 15 OS=Mus musculus OX=10090 GN=Rbm15 PE=1 SV=1 | 105.7 | 3 |
| Q8K019 | Bcl-2-associated transcription factor 1 OS=Mus musculus OX=10090 GN=Bclaf1 PE=1 SV=2 | 105.9 | 6 |
| Q569Z6 | Thyroid hormone receptor-associated protein 3 OS=Mus musculus OX=10090 GN=Thrap3 PE=1 SV=1 | 108.1 | 10 |

| | | | |
|---|---|---|---|
| Q3UMY5 | Echinoderm microtubule-associated protein-like 4 OS=Mus musculus OX=10090 GN=Eml4 PE=1 SV=1 | 110.0 | 3 |
| Q7TQH0 | Ataxin-2-like protein OS=Mus musculus OX=10090 GN=Atxn2l PE=1 SV=1 | 110.6 | 7 |
| O55098 | Serine/threonine-protein kinase 10 OS=Mus musculus OX=10090 GN=Stk10 PE=1 SV=2 | 111.8 | 3 |
| O35464 | Semaphorin-6A OS=Mus musculus OX=10090 GN=Sema6a PE=1 SV=2 | 114.4 | 2 |
| O09159 | Lysosomal alpha-mannosidase OS=Mus musculus OX=10090 GN=Man2b1 PE=1 SV=4 | 114.6 | 3 |
| Q9DBR7 | Protein phosphatase 1 regulatory subunit 12A OS=Mus musculus OX=10090 GN=Ppp1r12a PE=1 SV=2 | 114.9 | 2 |
| P27546 | Microtubule-associated protein 4 OS=Mus musculus OX=10090 GN=Map4 PE=1 SV=3 | 117.4 | 2 |
| Q91VX2 | Ubiquitin-associated protein 2 OS=Mus musculus OX=10090 GN=Ubap2 PE=1 SV=1 | 117.9 | 2 |
| P34152 | Focal adhesion kinase 1 OS=Mus musculus OX=10090 GN=Ptk2 PE=1 SV=4 | 119.2 | 7 |
| Q80U87 | Ubiquitin carboxyl-terminal hydrolase 8 OS=Mus musculus OX=10090 GN=Usp8 PE=1 SV=2 | 122.5 | 2 |
| P42703 | Leukemia inhibitory factor receptor OS=Mus musculus OX=10090 GN=Lifr PE=1 SV=1 | 122.5 | 3 |
| Q8CGF7 | Transcription elongation regulator 1 OS=Mus musculus OX=10090 GN=Tcerg1 PE=1 SV=2 | 123.7 | 2 |
| O08784 | Treacle protein OS=Mus musculus OX=10090 GN=Tcof1 PE=1 SV=1 | 134.9 | 2 |
| O70305 | Ataxin-2 OS=Mus musculus OX=10090 GN=Atxn2 PE=1 SV=1 | 136.4 | 4 |
| Q9EPX2 | Papilin OS=Mus musculus OX=10090 GN=Papln PE=2 SV=2 | 138.8 | 19 |
| Q6ZPZ3 | Zinc finger CCCH domain-containing protein 4 OS=Mus musculus OX=10090 GN=Zc3h4 PE=1 SV=2 | 140.9 | 3 |
| Q62417 | Sorbin and SH3 domain-containing protein 1 OS=Mus musculus OX=10090 GN=Sorbs1 PE=1 SV=2 | 143.0 | 3 |
| Q8K4P0 | pre-mRNA 3' end processing protein WDR33 OS=Mus musculus OX=10090 GN=Wdr33 PE=1 SV=1 | 145.2 | 2 |
| Q7TPV4 | Myb-binding protein 1A OS=Mus musculus OX=10090 GN=Mybbp1a PE=1 SV=2 | 151.9 | 4 |
| Q9ESU6 | Bromodomain-containing protein 4 OS=Mus musculus OX=10090 GN=Brd4 PE=1 SV=2 | 155.8 | 2 |
| P23116 | Eukaryotic translation initiation factor 3 subunit A OS=Mus musculus OX=10090 GN=Eif3a PE=1 SV=5 | 161.8 | 3 |
| Q8CGC7 | Bifunctional glutamate/proline--tRNA ligase OS=Mus musculus OX=10090 GN=Eprs1 PE=1 SV=4 | 170.0 | 7 |
| Q99PL5 | Ribosome-binding protein 1 OS=Mus musculus OX=10090 GN=Rrbp1 PE=1 SV=2 | 172.8 | 6 |
| B2RX14 | Terminal uridylyltransferase 4 OS=Mus musculus OX=10090 GN=Tut4 PE=1 SV=2 | 184.5 | 2 |
| Q9JKF1 | Ras GTPase-activating-like protein IQGAP1 OS=Mus musculus OX=10090 GN=Iqgap1 PE=1 SV=2 | 188.6 | 2 |
| P97868 | E3 ubiquitin-protein ligase RBBP6 OS=Mus musculus OX=10090 GN=Rbbp6 PE=1 SV=5 | 199.5 | 3 |
| B0V2N1 | Receptor-type tyrosine-protein phosphatase S OS=Mus musculus OX=10090 GN=Ptprs PE=1 SV=1 | 211.8 | 2 |
| F6ZDS4 | Nucleoprotein TPR OS=Mus musculus OX=10090 GN=Tpr PE=1 SV=1 | 273.8 | 5 |

| | | | |
|---|---|---|---|
| Q8BTM8 | Filamin-A OS=Mus musculus OX=10090 GN=Flna PE=1 SV=5 | 281.0 | 5 |
| Q3TLH4 | Protein PRRC2C OS=Mus musculus OX=10090 GN=Prrc2c PE=1 SV=3 | 310.7 | 8 |
| Q6KCD5 | Nipped-B-like protein OS=Mus musculus OX=10090 GN=Nipbl PE=1 SV=1 | 315.3 | 6 |
| Q62504 | Msx2-interacting protein OS=Mus musculus OX=10090 GN=Spen PE=1 SV=2 | 398.5 | 3 |
| Q8R0W0 | Epiplakin OS=Mus musculus OX=10090 GN=Eppk1 PE=1 SV=2 | 724.2 | 2 |

(**C**)

| Accession | Description | MW, kDa | # Unique Peptides |
|---|---|---|---|
| P62075 | Mitochondrial import inner membrane translocase subunit Tim13 OS=Mus musculus OX=10090 GN=Timm13 PE=1 SV=1 | 10.5 | 4 |
| Q64433 | 10 kDa heat shock protein, mitochondrial OS=Mus musculus OX=10090 GN=Hspe1 PE=1 SV=2 | 11.0 | 3 |
| Q9CWW6 | Peptidyl-prolyl cis-trans isomerase NIMA-interacting 4 OS=Mus musculus OX=10090 GN=Pin4 PE=1 SV=1 | 13.8 | 3 |
| Q9D8S9 | BolA-like protein 1 OS=Mus musculus OX=10090 GN=Bola1 PE=1 SV=1 | 14.4 | 2 |
| Q03958 | Prefoldin subunit 6 OS=Mus musculus OX=10090 GN=Pfdn6 PE=1 SV=1 | 14.4 | 6 |
| P59708 | Splicing factor 3B subunit 6 OS=Mus musculus OX=10090 GN=Sf3b6 PE=1 SV=1 | 14.6 | 2 |
| Q9CR98 | Protein FAM136A OS=Mus musculus OX=10090 GN=Fam136a PE=1 SV=1 | 15.7 | 2 |
| Q9CZG9 | PDZ domain-containing protein 11 OS=Mus musculus OX=10090 GN=Pdzd11 PE=1 SV=1 | 16.2 | 2 |
| O70591 | Prefoldin subunit 2 OS=Mus musculus OX=10090 GN=Pfdn2 PE=1 SV=2 | 16.5 | 2 |
| Q9CQ92 | Mitochondrial fission 1 protein OS=Mus musculus OX=10090 GN=Fis1 PE=1 SV=1 | 17.0 | 3 |
| P62843 | Small ribosomal subunit protein uS19 OS=Mus musculus OX=10090 GN=Rps15 PE=1 SV=2 | 17.0 | 3 |
| P61089 | Ubiquitin-conjugating enzyme E2 N OS=Mus musculus OX=10090 GN=Ube2n PE=1 SV=1 | 17.1 | 2 |
| P62270 | Small ribosomal subunit protein uS13 OS=Mus musculus OX=10090 GN=Rps18 PE=1 SV=3 | 17.7 | 4 |
| P21126 | Ubiquitin-like protein 4A OS=Mus musculus OX=10090 GN=Ubl4a PE=1 SV=1 | 17.8 | 2 |
| Q9DCT6 | Chromatin complexes subunit BAP18 OS=Mus musculus OX=10090 GN=Bap18 PE=1 SV=1 | 18.0 | 2 |
| Q9CXZ1 | NADH dehydrogenase [ubiquinone] iron-sulfur protein 4, mitochondrial OS=Mus musculus OX=10090 GN=Ndufs4 PE=1 SV=3 | 19.8 | 2 |
| Q6PGH2 | Jupiter microtubule associated homolog 2 OS=Mus musculus OX=10090 GN=Jpt2 PE=1 SV=1 | 20.0 | 3 |
| Q99LX0 | Parkinson disease protein 7 homolog OS=Mus musculus OX=10090 GN=Park7 PE=1 SV=1 | 20.0 | 5 |
| P61205 | ADP-ribosylation factor 3 OS=Mus musculus OX=10090 GN=Arf3 PE=2 SV=2 | 20.6 | 3 |
| Q61171 | Peroxiredoxin-2 OS=Mus musculus OX=10090 GN=Prdx2 PE=1 SV=3 | 21.8 | 3 |
| Q9CQW1 | Synaptobrevin homolog YKT6 OS=Mus musculus OX=10090 GN=Ykt6 PE=1 SV=1 | 22.3 | 3 |
| Q8VCG4 | Complement component C8 gamma chain OS=Mus musculus OX=10090 GN=C8g PE=1 SV=1 | 22.5 | 4 |
| Q8BPA8 | Protein DPCD OS=Mus musculus OX=10090 GN=Dpcd PE=1 SV=1 | 23.0 | 6 |
| Q78PG9 | Coiled-coil domain-containing protein 25 OS=Mus musculus OX=10090 GN=Ccdc25 PE=1 SV=1 | 24.5 | 3 |
| P09671 | Superoxide dismutase [Mn], mitochondrial OS=Mus musculus OX=10090 GN=Sod2 PE=1 SV=3 | 24.6 | 5 |

| | | | |
|---|---|---|---|
| P63158 | High mobility group protein B1 OS=Mus musculus OX=10090 GN=Hmgb1 PE=1 SV=2 | 24.9 | 3 |
| Q9ERE7 | LRP chaperone MESD OS=Mus musculus OX=10090 GN=Mesd PE=1 SV=1 | 25.2 | 2 |
| P10649 | Glutathione S-transferase Mu 1 OS=Mus musculus OX=10090 GN=Gstm1 PE=1 SV=2 | 26.0 | 2 |
| Q9CRB9 | MICOS complex subunit Mic19 OS=Mus musculus OX=10090 GN=Chchd3 PE=1 SV=1 | 26.3 | 2 |
| Q8C7V8 | Coiled-coil domain-containing protein 134 OS=Mus musculus OX=10090 GN=Ccdc134 PE=1 SV=1 | 26.5 | 3 |
| P17751 | Triosephosphate isomerase OS=Mus musculus OX=10090 GN=Tpi1 PE=1 SV=5 | 26.7 | 2 |
| O88983 | Syntaxin-8 OS=Mus musculus OX=10090 GN=Stx8 PE=1 SV=1 | 26.9 | 2 |
| O09131 | Glutathione S-transferase omega-1 OS=Mus musculus OX=10090 GN=Gsto1 PE=1 SV=2 | 27.5 | 5 |
| Q9CQE8 | RNA transcription, translation and transport factor protein OS=Mus musculus OX=10090 GN=RTRAF PE=1 SV=1 | 28.1 | 4 |
| Q9D172 | Glutamine amidotransferase-like class 1 domain-containing protein 3, mitochondrial OS=Mus musculus OX=10090 GN=Gatd3 PE=1 SV=1 | 28.1 | 4 |
| Q9D1J1 | Adaptin ear-binding coat-associated protein 2 OS=Mus musculus OX=10090 GN=Necap2 PE=1 SV=1 | 28.6 | 2 |
| O70250 | Phosphoglycerate mutase 2 OS=Mus musculus OX=10090 GN=Pgam2 PE=1 SV=3 | 28.8 | 4 |
| P62259 | 14-3-3 protein epsilon OS=Mus musculus OX=10090 GN=Ywhae PE=1 SV=1 | 29.2 | 2 |
| Q9QZH3 | Peptidyl-prolyl cis-trans isomerase E OS=Mus musculus OX=10090 GN=Ppie PE=1 SV=2 | 33.4 | 2 |
| Q922Y1 | UBX domain-containing protein 1 OS=Mus musculus OX=10090 GN=Ubxn1 PE=1 SV=1 | 33.6 | 4 |
| P10711 | Transcription elongation factor A protein 1 OS=Mus musculus OX=10090 GN=Tcea1 PE=1 SV=2 | 33.9 | 2 |
| Q99KB8 | Hydroxyacylglutathione hydrolase, mitochondrial OS=Mus musculus OX=10090 GN=Hagh PE=1 SV=2 | 34.1 | 2 |
| Q9Z204 | Heterogeneous nuclear ribonucleoproteins C1/C2 OS=Mus musculus OX=10090 GN=Hnrnpc PE=1 SV=1 | 34.4 | 5 |
| Q61425 | Hydroxyacyl-coenzyme A dehydrogenase, mitochondrial OS=Mus musculus OX=10090 GN=Hadh PE=1 SV=2 | 34.4 | 5 |
| P56528 | ADP-ribosyl cyclase/cyclic ADP-ribose hydrolase 1 OS=Mus musculus OX=10090 GN=Cd38 PE=1 SV=2 | 34.4 | 2 |
| Q8BM88 | Cathepsin O OS=Mus musculus OX=10090 GN=Ctso PE=2 SV=1 | 34.7 | 2 |
| Q61176 | Arginase-1 OS=Mus musculus OX=10090 GN=Arg1 PE=1 SV=1 | 34.8 | 4 |
| Q91Z53 | Glyoxylate reductase/hydroxypyruvate reductase OS=Mus musculus OX=10090 GN=Grhpr PE=1 SV=1 | 35.3 | 2 |
| P45376 | Aldo-keto reductase family 1 member B1 OS=Mus musculus OX=10090 GN=Akr1b1 PE=1 SV=3 | 35.7 | 5 |
| P62960 | Y-box-binding protein 1 OS=Mus musculus OX=10090 GN=Ybx1 PE=1 SV=3 | 35.7 | 3 |
| P16858 | Glyceraldehyde-3-phosphate dehydrogenase OS=Mus musculus OX=10090 GN=Gapdh PE=1 SV=2 | 35.8 | 2 |
| P14152 | Malate dehydrogenase, cytoplasmic OS=Mus musculus OX=10090 GN=Mdh1 PE=1 SV=3 | 36.5 | 2 |

| | | | |
|---|---|---|---|
| Q9D1P4 | Cysteine and histidine-rich domain-containing protein 1 OS=Mus musculus OX=10090 GN=Chordc1 PE=1 SV=1 | 37.3 | 2 |
| Q9ES89 | Exostosin-like 2 OS=Mus musculus OX=10090 GN=Extl2 PE=1 SV=1 | 37.4 | 6 |
| Q8VCN9 | Tubulin-specific chaperone C OS=Mus musculus OX=10090 GN=Tbcc PE=1 SV=1 | 38.1 | 2 |
| Q64442 | Sorbitol dehydrogenase OS=Mus musculus OX=10090 GN=Sord PE=1 SV=3 | 38.2 | 6 |
| O35685 | Nuclear migration protein nudC OS=Mus musculus OX=10090 GN=Nudc PE=1 SV=1 | 38.3 | 2 |
| P24452 | Macrophage-capping protein OS=Mus musculus OX=10090 GN=Capg PE=1 SV=2 | 39.2 | 4 |
| Q3UMW8 | Bis(monoacylglycero)phosphate synthase CLN5 OS=Mus musculus OX=10090 GN=Cln5 PE=1 SV=1 | 39.3 | 3 |
| Q9JHJ0 | Tropomodulin-3 OS=Mus musculus OX=10090 GN=Tmod3 PE=1 SV=1 | 39.5 | 2 |
| Q8BG05 | Heterogeneous nuclear ribonucleoprotein A3 OS=Mus musculus OX=10090 GN=Hnrnpa3 PE=1 SV=1 | 39.6 | 2 |
| P54726 | UV excision repair protein RAD23 homolog A OS=Mus musculus OX=10090 GN=Rad23a PE=1 SV=2 | 39.7 | 2 |
| O54946 | DnaJ homolog subfamily B member 6 OS=Mus musculus OX=10090 GN=Dnajb6 PE=1 SV=4 | 39.8 | 4 |
| Q9CX56 | 26S proteasome non-ATPase regulatory subunit 8 OS=Mus musculus OX=10090 GN=Psmd8 PE=1 SV=2 | 39.9 | 3 |
| Q00731-6 | Isoform L-VEGF-1 of Vascular endothelial growth factor A, long form OS=Mus musculus OX=10090 GN=Vegfa | 40.3 | 2 |
| P63085 | Mitogen-activated protein kinase 1 OS=Mus musculus OX=10090 GN=Mapk1 PE=1 SV=3 | 41.2 | 4 |
| Q8CAY6 | Acetyl-CoA acetyltransferase, cytosolic OS=Mus musculus OX=10090 GN=Acat2 PE=1 SV=2 | 41.3 | 4 |
| Q9Z0P4 | Paralemmin-1 OS=Mus musculus OX=10090 GN=Palm PE=1 SV=1 | 41.6 | 2 |
| P50580 | Proliferation-associated protein 2G4 OS=Mus musculus OX=10090 GN=Pa2g4 PE=1 SV=3 | 43.7 | 2 |
| Q61187 | Tumor susceptibility gene 101 protein OS=Mus musculus OX=10090 GN=Tsg101 PE=1 SV=2 | 44.1 | 2 |
| O89112 | Glutathione S-transferase LANCL1 OS=Mus musculus OX=10090 GN=Lancl1 PE=1 SV=1 | 45.3 | 3 |
| Q8BK62 | Olfactomedin-like protein 3 OS=Mus musculus OX=10090 GN=Olfml3 PE=2 SV=2 | 45.7 | 2 |
| Q8BV49 | Pyrin and HIN domain-containing protein 1 OS=Mus musculus OX=10090 GN=Pyhin1 PE=1 SV=1 | 46.9 | 2 |
| Q8BHS3 | Pre-mRNA-splicing factor RBM22 OS=Mus musculus OX=10090 GN=Rbm22 PE=1 SV=1 | 46.9 | 2 |
| Q9QWR8 | Alpha-N-acetylgalactosaminidase OS=Mus musculus OX=10090 GN=Naga PE=1 SV=2 | 47.2 | 2 |
| Q8BP40 | Lysophosphatidic acid phosphatase type 6 OS=Mus musculus OX=10090 GN=Acp6 PE=1 SV=1 | 47.6 | 3 |
| P50247 | Adenosylhomocysteinase OS=Mus musculus OX=10090 GN=Ahcy PE=1 SV=3 | 47.7 | 9 |
| Q8VEJ9 | Vacuolar protein sorting-associated protein 4A OS=Mus musculus OX=10090 GN=Vps4a PE=1 SV=1 | 48.9 | 2 |
| Q9JIY5 | Serine protease HTRA2, mitochondrial OS=Mus musculus OX=10090 GN=Htra2 PE=1 SV=2 | 49.3 | 5 |

| | | | |
|---|---|---|---|
| P61979 | Heterogeneous nuclear ribonucleoprotein K OS=Mus musculus OX=10090 GN=Hnrnpk PE=1 SV=1 | 50.9 | 4 |
| P27808 | Alpha-1,3-mannosyl-glycoprotein 2-beta-N-acetylglucosaminyltransferase OS=Mus musculus OX=10090 GN=Mgat1 PE=1 SV=1 | 51.7 | 2 |
| Q9Z2W0 | Aspartyl aminopeptidase OS=Mus musculus OX=10090 GN=Dnpep PE=1 SV=2 | 52.2 | 3 |
| Q9DCD0 | 6-phosphogluconate dehydrogenase, decarboxylating OS=Mus musculus OX=10090 GN=Pgd PE=1 SV=3 | 53.2 | 8 |
| Q56A08 | G-patch domain and KOW motifs-containing protein OS=Mus musculus OX=10090 GN=Gpkow PE=1 SV=2 | 53.8 | 3 |
| Q2TPA8 | Hydroxysteroid dehydrogenase-like protein 2 OS=Mus musculus OX=10090 GN=Hsdl2 PE=1 SV=1 | 54.2 | 2 |
| Q8BG07 | 5'-3' exonuclease PLD4 OS=Mus musculus OX=10090 GN=Pld4 PE=1 SV=1 | 56.1 | 5 |
| Q61753 | D-3-phosphoglycerate dehydrogenase OS=Mus musculus OX=10090 GN=Phgdh PE=1 SV=3 | 56.5 | 4 |
| Q99K28 | ADP-ribosylation factor GTPase-activating protein 2 OS=Mus musculus OX=10090 GN=Arfgap2 PE=1 SV=1 | 56.6 | 5 |
| Q9D8S3 | ADP-ribosylation factor GTPase-activating protein 3 OS=Mus musculus OX=10090 GN=Arfgap3 PE=1 SV=2 | 57.4 | 5 |
| Q8BG30 | Negative elongation factor A OS=Mus musculus OX=10090 GN=Nelfa PE=1 SV=1 | 57.5 | 3 |
| P52480 | Pyruvate kinase PKM OS=Mus musculus OX=10090 GN=Pkm PE=1 SV=4 | 57.8 | 5 |
| O08795 | Glucosidase 2 subunit beta OS=Mus musculus OX=10090 GN=Prkcsh PE=1 SV=1 | 58.8 | 2 |
| P63038 | 60 kDa heat shock protein, mitochondrial OS=Mus musculus OX=10090 GN=Hspd1 PE=1 SV=1 | 60.9 | 4 |
| Q8BFR4 | N-acetylglucosamine-6-sulfatase OS=Mus musculus OX=10090 GN=Gns PE=1 SV=1 | 61.1 | 2 |
| Q8BIQ5 | Cleavage stimulation factor subunit 2 OS=Mus musculus OX=10090 GN=Cstf2 PE=1 SV=2 | 61.3 | 2 |
| Q61712 | DnaJ homolog subfamily C member 1 OS=Mus musculus OX=10090 GN=Dnajc1 PE=1 SV=1 | 63.8 | 3 |
| Q925I1 | ATPase family AAA domain-containing protein 3 OS=Mus musculus OX=10090 GN=Atad3 PE=1 SV=1 | 66.7 | 2 |
| Q8CHY6 | Transcriptional repressor p66 alpha OS=Mus musculus OX=10090 GN=Gatad2a PE=1 SV=2 | 67.3 | 2 |
| P35235-1 | Isoform 2 of Tyrosine-protein phosphatase non-receptor type 11 OS=Mus musculus OX=10090 GN=Ptpn11 | 68.4 | 7 |
| Q91WJ8 | Far upstream element-binding protein 1 OS=Mus musculus OX=10090 GN=Fubp1 PE=1 SV=1 | 68.5 | 5 |
| P10404 | MLV-related proviral Env polyprotein OS=Mus musculus OX=10090 PE=1 SV=3 | 69.6 | 2 |
| Q9JL61 | DNA-binding protein Rfx5 OS=Mus musculus OX=10090 GN=Rfx5 PE=1 SV=2 | 69.7 | 2 |
| Q8R0H9 | ADP-ribosylation factor-binding protein GGA1 OS=Mus musculus OX=10090 GN=Gga1 PE=1 SV=1 | 69.9 | 2 |
| P54729 | NEDD8 ultimate buster 1 OS=Mus musculus OX=10090 GN=Nub1 PE=1 SV=2 | 70.3 | 2 |
| Q80TY0 | Formin-binding protein 1 OS=Mus musculus OX=10090 GN=Fnbp1 PE=1 SV=2 | 71.3 | 3 |
| P08003 | Protein disulfide-isomerase A4 OS=Mus musculus OX=10090 GN=Pdia4 PE=1 SV=3 | 71.9 | 3 |

| | | | |
|---|---|---|---|
| Q8BVL9 | Janus kinase and microtubule-interacting protein 1 OS=Mus musculus OX=10090 GN=Jakmip1 PE=1 SV=2 | 73.1 | 2 |
| Q8VC60 | Beta-galactosidase-1-like protein OS=Mus musculus OX=10090 GN=Glb1l PE=1 SV=1 | 73.2 | 5 |
| Q3V1H1 | Cytoskeleton-associated protein 2 OS=Mus musculus OX=10090 GN=Ckap2 PE=1 SV=1 | 74.0 | 2 |
| Q5F2E7 | FMR1-interacting protein NUFIP2 OS=Mus musculus OX=10090 GN=Nufip2 PE=1 SV=1 | 75.6 | 2 |
| Q3U0V1 | Far upstream element-binding protein 2 OS=Mus musculus OX=10090 GN=Khsrp PE=1 SV=2 | 76.7 | 6 |
| Q9D0E1 | Heterogeneous nuclear ribonucleoprotein M OS=Mus musculus OX=10090 GN=Hnrnpm PE=1 SV=3 | 77.6 | 4 |
| P51660 | Peroxisomal multifunctional enzyme type 2 OS=Mus musculus OX=10090 GN=Hsd17b4 PE=1 SV=3 | 79.4 | 2 |
| Q08943 | FACT complex subunit SSRP1 OS=Mus musculus OX=10090 GN=Ssrp1 PE=1 SV=2 | 80.8 | 2 |
| Q9CZD3 | Glycine--tRNA ligase OS=Mus musculus OX=10090 GN=Gars1 PE=1 SV=1 | 81.8 | 2 |
| Q61183 | Poly(A) polymerase alpha OS=Mus musculus OX=10090 GN=Papola PE=1 SV=4 | 82.3 | 2 |
| Q8BJ05 | Zinc finger CCCH domain-containing protein 14 OS=Mus musculus OX=10090 GN=Zc3h14 PE=1 SV=1 | 82.4 | 4 |
| Q3UIR3 | E3 ubiquitin-protein ligase DTX3L OS=Mus musculus OX=10090 GN=Dtx3l PE=1 SV=1 | 83.0 | 6 |
| Q99KI0 | Aconitate hydratase, mitochondrial OS=Mus musculus OX=10090 GN=Aco2 PE=1 SV=1 | 85.4 | 17 |
| Q9QYH6 | Melanoma-associated antigen D1 OS=Mus musculus OX=10090 GN=Maged1 PE=1 SV=1 | 85.6 | 2 |
| Q62351 | Transferrin receptor protein 1 OS=Mus musculus OX=10090 GN=Tfrc PE=1 SV=1 | 85.7 | 3 |
| Q99LI8 | Hepatocyte growth factor-regulated tyrosine kinase substrate OS=Mus musculus OX=10090 GN=Hgs PE=1 SV=2 | 86.0 | 3 |
| Q6NZF1 | Zinc finger CCCH domain-containing protein 11A OS=Mus musculus OX=10090 GN=Zc3h11a PE=1 SV=1 | 86.4 | 2 |
| Q924H2 | Mediator of RNA polymerase II transcription subunit 15 OS=Mus musculus OX=10090 GN=Med15 PE=1 SV=3 | 86.6 | 3 |
| Q922K7 | 28S rRNA (cytosine-C(5))-methyltransferase OS=Mus musculus OX=10090 GN=Nop2 PE=1 SV=1 | 86.7 | 2 |
| P27612 | Phospholipase A-2-activating protein OS=Mus musculus OX=10090 GN=Plaa PE=1 SV=4 | 87.2 | 2 |
| Q8BML9 | Glutamine--tRNA ligase OS=Mus musculus OX=10090 GN=Qars1 PE=1 SV=1 | 87.6 | 3 |
| P25976 | Nucleolar transcription factor 1 OS=Mus musculus OX=10090 GN=Ubtf PE=1 SV=1 | 89.5 | 2 |
| Q00547 | Hyaluronan mediated motility receptor OS=Mus musculus OX=10090 GN=Hmmr PE=1 SV=4 | 91.7 | 2 |
| Q9WUH7 | Semaphorin-4G OS=Mus musculus OX=10090 GN=Sema4g PE=1 SV=1 | 92.3 | 5 |
| P08113 | Endoplasmin OS=Mus musculus OX=10090 GN=Hsp90b1 PE=1 SV=2 | 92.4 | 2 |
| Q61316 | Heat shock 70 kDa protein 4 OS=Mus musculus OX=10090 GN=Hspa4 PE=1 SV=1 | 94.1 | 2 |
| P42567 | Epidermal growth factor receptor substrate 15 OS=Mus musculus OX=10090 GN=Eps15 PE=1 SV=1 | 98.4 | 2 |

| | | | |
|---|---|---|---|
| Q9R1E6 | Autotaxin OS=Mus musculus OX=10090 GN=Enpp2 PE=1 SV=3 | 98.8 | 3 |
| B2RY56 | RNA-binding protein 25 OS=Mus musculus OX=10090 GN=Rbm25 PE=1 SV=2 | 99.5 | 2 |
| P56960 | Exosome complex component 10 OS=Mus musculus OX=10090 GN=Exosc10 PE=1 SV=2 | 100.9 | 2 |
| Q52KI8 | Serine/arginine repetitive matrix protein 1 OS=Mus musculus OX=10090 GN=Srrm1 PE=1 SV=2 | 106.8 | 3 |
| A2A432 | Cullin-4B OS=Mus musculus OX=10090 GN=Cul4b PE=1 SV=1 | 110.6 | 3 |
| P11103 | Poly [ADP-ribose] polymerase 1 OS=Mus musculus OX=10090 GN=Parp1 PE=1 SV=3 | 113.0 | 4 |
| Q80X50 | Ubiquitin-associated protein 2-like OS=Mus musculus OX=10090 GN=Ubap2l PE=1 SV=1 | 116.7 | 3 |
| Q569Z5 | Probable ATP-dependent RNA helicase DDX46 OS=Mus musculus OX=10090 GN=Ddx46 PE=1 SV=2 | 117.4 | 4 |
| Q920B9 | FACT complex subunit SPT16 OS=Mus musculus OX=10090 GN=Supt16h PE=1 SV=2 | 119.7 | 3 |
| O55201 | Transcription elongation factor SPT5 OS=Mus musculus OX=10090 GN=Supt5h PE=1 SV=1 | 120.6 | 2 |
| Q61147 | Ceruloplasmin OS=Mus musculus OX=10090 GN=Cp PE=1 SV=2 | 121.1 | 4 |
| Q05D44 | Eukaryotic translation initiation factor 5B OS=Mus musculus OX=10090 GN=Eif5b PE=1 SV=2 | 137.5 | 2 |
| P33174 | Chromosome-associated kinesin KIF4 OS=Mus musculus OX=10090 GN=Kif4 PE=1 SV=3 | 139.4 | 3 |
| Q99KY4 | Cyclin-G-associated kinase OS=Mus musculus OX=10090 GN=Gak PE=1 SV=2 | 143.6 | 3 |
| Q8CG47 | Structural maintenance of chromosomes protein 4 OS=Mus musculus OX=10090 GN=Smc4 PE=1 SV=1 | 146.8 | 4 |
| Q9JIX8 | Apoptotic chromatin condensation inducer in the nucleus OS=Mus musculus OX=10090 GN=Acin1 PE=1 SV=3 | 150.6 | 4 |
| Q61595 | Kinectin OS=Mus musculus OX=10090 GN=Ktn1 PE=1 SV=1 | 152.5 | 2 |
| Q5FWI3 | Cell surface hyaluronidase OS=Mus musculus OX=10090 GN=Cemip2 PE=1 SV=1 | 153.7 | 2 |
| Q922J3 | CAP-Gly domain-containing linker protein 1 OS=Mus musculus OX=10090 GN=Clip1 PE=1 SV=1 | 155.7 | 3 |
| Q9EQJ9 | Membrane-associated guanylate kinase, WW and PDZ domain-containing protein 3 OS=Mus musculus OX=10090 GN=Magi3 PE=1 SV=2 | 161.6 | 2 |
| P13864 | DNA (cytosine-5)-methyltransferase 1 OS=Mus musculus OX=10090 GN=Dnmt1 PE=1 SV=5 | 183.1 | 4 |
| Q9Z0R4 | Intersectin-1 OS=Mus musculus OX=10090 GN=Itsn1 PE=1 SV=2 | 194.2 | 4 |
| A2ASQ1 | Agrin OS=Mus musculus OX=10090 GN=Agrn PE=1 SV=1 | 207.4 | 2 |
| Q80UG2 | Plexin-A4 OS=Mus musculus OX=10090 GN=Plxna4 PE=1 SV=3 | 212.4 | 6 |
| Q8VDD5 | Myosin-9 OS=Mus musculus OX=10090 GN=Myh9 PE=1 SV=4 | 226.2 | 3 |
| B2RWS6 | Histone acetyltransferase p300 OS=Mus musculus OX=10090 GN=Ep300 PE=1 SV=2 | 263.1 | 2 |
| Q62261 | Spectrin beta chain, non-erythrocytic 1 OS=Mus musculus OX=10090 GN=Sptbn1 PE=1 SV=2 | 274.1 | 2 |
| Q811L6 | Microtubule-associated serine/threonine-protein kinase 4 OS=Mus musculus OX=10090 GN=Mast4 PE=1 SV=3 | 283.8 | 6 |

| | | | |
|---|---|---|---|
| Q8BTI8 | Serine/arginine repetitive matrix protein 2 OS=Mus musculus OX=10090 GN=Srrm2 PE=1 SV=3 | 294.7 | 3 |
| Q9QYR6 | Microtubule-associated protein 1A OS=Mus musculus OX=10090 GN=Map1a PE=1 SV=2 | 300.0 | 2 |
| E9Q6J5 | Biorientation of chromosomes in cell division protein 1-like 1 OS=Mus musculus OX=10090 GN=Bod1l PE=1 SV=1 | 327.3 | 2 |
| E9PVX6 | Proliferation marker protein Ki-67 OS=Mus musculus OX=10090 GN=Mki67 PE=1 SV=1 | 350.7 | 2 |
